# The Interplay Between Multisensory Processing and Attention in Working Memory: Behavioral and Neural Indices of Audio-Visual Object Storage

**DOI:** 10.1101/2024.03.26.586805

**Authors:** Ceren Arslan, Daniel Schneider, Stephan Getzmann, Edmund Wascher, Laura-Isabelle Klatt

**Affiliations:** Leibniz Research Centre for Working Environment and Human Factors

**Keywords:** selective attention, working memory, multisensory processing, alpha oscillations, EEG

## Abstract

Although real-life events are multisensory, how audio-visual objects are stored in working memory is an open question. At a perceptual level, evidence shows that both top-down and bottom-up attentional processes can play a role in multisensory interactions. To understand how attention and multisensory processes interact in working memory, we designed an audio-visual delayed match-to-sample task in which participants were presented with one or two audio-visual memory items, followed by an audio-visual probe. In three different blocks, participants were instructed to either (a) attend to the auditory features, (b) attend to the visual features, or (c) attend to both auditory and visual features. Participants were instructed to indicate whether the task-relevant features of the probe matched one of the task-relevant feature(s) or objects in working memory. Behavioral results showed interference from task-irrelevant features, suggesting bottom-up integration of audio-visual features and their automatic encoding into working memory, irrespective of their task relevance. Yet, ERP analyses revealed no evidence for active maintenance of these task-irrelevant features, while they clearly taxed greater attentional resources during recall. Notably, alpha oscillatory activity revealed that linking information between auditory and visual modalities required more attentional demands at retrieval. Overall, these results offer critical insights into how and at which processing stage multisensory interactions occur in working memory.

**Public Significance Statement:** Current working memory research is dominated by investigations of the visual domain. Yet, understanding how more complex representations, e.g., based on multisensory inputs, are formed, stored, and recalled is crucial to obtaining a more realistic understanding of working memory function. The present study shows that when confronted with audio-visual inputs at the same time and location, features from both modalities are combined into a working memory representation, irrespective of their task relevance. During maintenance, alpha oscillations serve to flexibly gate information flow in the cortex, allowing for the re-distribution of attentional resources between modalities depending on their task-relevance. Notably, when the task instructions explicitly involve the storage of audio-visual objects as a whole, recall requires more attentional resources.

## Introduction

To navigate through a complex, multisensory environment, our senses need to interact with each other constantly. This applies not only to low-level perceptual mechanisms but also to higher-order cognitive functions such as working memory. However, current working memory research is strongly dominated by vision, and our understanding of how multisensory objects are encoded and maintained in working memory remains limited. The current study sets out to investigate how audio-visual information is stored in working memory and aims to observe neural correlates of attentional mechanisms supporting the linking of cross-modal information.

The concurrent storage of auditory and visual items has been predominantly investigated in a line of behavioral studies, adopting a dual-task paradigm (Cowan et al., 2014; Fougnie et al., 2015; Morey & Cowan, 2004, 2005; Saults & Cowan, 2007). Such studies primarily address the question as to whether storage limits in working memory are restricted by a central, amodal capacity or governed by domain-independent storage systems – a question that remains controversially debated and is beyond the scope of this article. Notably, these studies aim to isolate the subcomponents of working memory, as proposed by the classical modular view (Baddeley, 1986), rather than investigating multisensory working memory representations. Specifically, in most studies, the unimodal memory arrays are presented in spatiotemporal asynchrony, or only one of the modalities is probed during recall. Consequentially, previous studies did not create an incentive to encode or store cross-modally presented features as integrated audio-visual objects.

The question of how audio-visual information is encoded and maintained in working memory, ultimately also taps into the debate on whether individual features are stored separately or integrated as a bound object representation and to what extent the latter include task-irrelevant features. With respect to the perceptual aspects of cross-modal binding, we can draw on an ample body of studies in the multisensory processing literature, showing that the combination of information from different sensory modalities into a single multisensory event or representation (i.e., multisensory integration) can occur in a purely bottom-up manner if certain conditions are met, but requires an additional top-down attention mechanism otherwise (Busse et al., 2005; Matusz et al., 2019; Sarmiento et al., 2016; Senkowski et al., 2005, 2007, 2009). According to a framework by Talsma and colleagues (2010), the interplay between attention and multisensory processing is determined by the physical complexity of the environment. If the competition for a limited capacity between stimuli within a given modality is low, cross-modal interactions can happen in a bottom-up fashion. Accordingly, the co-occurrence of auditory and visual stimuli in space and time is likely to promote the cross-modal spread of attention towards task-irrelevant features of an attended object. Supporting this idea, it has been shown that a task-irrelevant and spatially uninformative sound that is presented concurrently with a change of a visual target item facilitates visual search task performance (Van Der Burg et al., 2008, 2010). This observation has been interpreted as evidence for the bottom-up driven integration of the task-irrelevant sound and the visual target, making the visual target pop-out from the cluttered scene.

Critically, whether incidentally or intentionally formed cross-modal feature bindings are also maintained in an integrated manner in working memory or whether the task-irrelevant feature is dropped from working memory, remains unclear. For visual working memory, a growing body of evidence suggests that object representations retain both a feature-as well as an object-level (Brady et al., 2011; Li et al., 2022). However, findings concerning the storage of task-irrelevant features remain somewhat contradictory. While some authors were able to successfully decode task-irrelevant features based on multivariate EEG data (Chen et al., 2021), others were only able to decode task-relevant features (Bocincova & Johnson, 2019; Yu & Shim, 2017). On the contrary, behavioral findings, such as inter-trial priming (Jiang et al., 2016) and attentional capture in a secondary search task (Harrison et al., 2021) corroborate the idea that at least some residual representation of task-irrelevant visual features is retained. Likewise, for auditory working memory, there is behavioral evidence showing better recall performance when participants are required to recall an auditory object as a whole compared to when they are asked to report individual features of the object, suggesting that auditory features might be naturally stored as bound objects (Joseph et al., 2015). Accordingly, Bays and colleagues (2022) postulated that task-irrelevant features of attended objects are automatically encoded, but not actively or only weakly maintained through the engagement of top-down resources. Yet, to date, with respect to multisensory object representations, this question remains largely unaddressed.

Although the representational format of multisensory information in working memory is not clear to this date, multiple working memory studies combining modality-specific or cross-modal stimulus presentation with electroencephalography (EEG), or magnetoencephalography (MEG) have illuminated the role of oscillatory mechanisms from various angles. Specifically, three major lines of research can be identified: One area of investigation has focused on identifying modality-independent, supramodal substrates of working memory retention or working memory control (Cowan et al., 2011; Majerus et al., 2016; Spitzer & Blankenburg, 2012; van Ede et al., 2017). In addition, several studies suggest that functional connectivity dynamics, supporting long-range communication between distant brain areas, may play an important role in the encoding of audio-visual objects (Xie et al., 2021), cross-modal integration (van Driel et al., 2014), as well as cognitive control of audio-visual working memory maintenance (Daume et al., 2017). Finally, a large body of literature delineates the key role of alpha (8-13 Hz) power modulations in facilitating unisensory (Dubé et al., 2013; Payne et al., 2013) as well as cross-modal (Foxe et al., 1998; Mazaheri et al., 2014) information processing by gating information between sensory regions (for a review see Jensen & Mazaheri, 2010). Extending this work from the perceptual domain, several working memory studies have demonstrated that alpha power likewise increases over mnemonically irrelevant sensory regions, while decreasing over task-relevant regions (Spitzer & Blankenburg, 2012; van Ede et al., 2017), corroborating its functional significance in the dynamic (dis-)engagement and functional inhibition of sensory areas.

Here, we recorded the EEG while participants completed three variants of an audio-visual delayed match-to-sample task. In two single-feature conditions, participants were instructed to attend to either the visual or the auditory features of the presented audio-visual objects, while in the conjunction condition (i.e., audio-visual condition) participants attended to both features. Compared to previous behavioral and EEG studies, the present design entails several key advances. Foremost, by always presenting an audio-visual (rather than a unisensory) probe, we prevent that the task itself entails an incentive to store auditory and visual features in a segregated manner. Further, by presenting the auditory and visual features at the same time and location, we promote the bottom-up integration of cross-modal input and thereby enhance the perceptual impression that both features originate from the same source and belong to the same multisensory object. Finally, we introduce a novel conjunction condition, in which participants are explicitly instructed to consider the auditory and visual features as part of the same object. Thereby, we create three variants of the task that are identical in terms of their physical stimulation (i.e., always present audio-visual input), but differ in terms of their task instruction.

The present study aims to address the following major questions: First, by assessing behavioral interference effects of the task-irrelevant features in the single-feature conditions, we aim to explore to what extend task-irrelevant cross-modal features are maintained in working memory. If the task-irrelevant features of memory items are stored to some degree, then they should interfere with the task performance; decreasing the accuracy and/or increasing reaction times when their comparison with the probe would imply a different response than the task-relevant feature. In addition, by exploiting well-established event-related potentials (ERPs) of auditory and visual working memory storage, we aim to track whether those task-irrelevant features are actively maintained. Such that, if task-uninformative auditory features are maintained in the visual condition, sustained anterior negativity amplitudes (i.e., SAN, an ERP correlate for auditory working memory storage, Alunni-Menichini et al., 2014; Guimond et al., 2011; Lefebvre et al., 2013; Nolden et al., 2013) should vary as the number of auditory features increases. Similarly, if visual features are actively maintained in the auditory condition, an analogous load effect should be visible in posterior negative slow-wave amplitudes (i.e., NSW, an ERP correlate for visual working memory storage, Diaz et al., 2021; Feldmann-Wüstefeld, 2021) as visual features increase in number. Further, while previous EEG studies have primarily focused on revealing the neurocognitive mechanisms supporting the prioritization of one modality over the other, the present study illuminates the oscillatory mechanisms involved in linking information across modalities by contrasting the single-feature conditions with the novel cross-modal conjunction condition. By inspecting alpha power modulations, the current study aims to observe the (re-)distribution of attentional resources when unimodal and cross-modal information is encoded, maintained and retrieved.

## Materials and Method

### Ethics statement

This study was approved by the Ethical Committee at the Leibniz Research Centre for Working Environment and Human Factors. It was carried out in line with the guidelines of the Declaration of Helsinki. All participants provided written informed consent before the experimental session. We compensated the participants with either 12 euros per hour or, if requested, course credits.

### Transparency and Openness

We discuss how we determined the sample size, exclusion criteria and manipulations in the study. The preregistration of the sample size rationale, experimental design, data analysis plan, and hypotheses of the study can be found at this link (https://osf.io/c5vjh/). Upon acceptance for publication, all research materials and data associated with this work will be publicly available in the respective link. Data were analyzed using JASP (version 0. 16.4.0) and MATLAB® (R2022a).

### Participants

50 participants took part in the study. Two participants were excluded from the dataset due to a disconnected ground electrode during EEG recording. Three participants were discarded from the dataset due to behavioral performance below or close to chance level (19%, 34%, and 56%, respectively). Finally, one participant was excluded due to a lack of cooperation in following the task instructions. The final sample included 44 participants (15 men, 29 women, and three left-handed) with a mean age of 23.18 years (*SD* = 3.59, age range = 18 to 34). This corresponds to the targeted sample size that was determined using an a-priori power analysis (see section ‘Power analysis’). All participants reported no history of neurological disorders or psychopharmacological medication.

We tested the hearing acuity of each participant using a pure-tone audiometry (Oscilla USB 330; Inmedico, Lystrup, Denmark), presenting eleven pure-tone frequencies in between 125 Hz and 8000 Hz. All participants showed normal hearing levels below or equal to 25 dB for frequencies between 125 Hz and 4 kHz. For frequencies higher than 4 kHz (i.e., 6 kHz and 8 kHz), five participants displayed marginally elevated hearing levels of 30 dB (3 participants at 6 kHz and 1 participant at 8 kHz) and 35 dB (1 participant at 6 kHz). Considering that all experimental auditory stimuli presented with a frequency below 4 kHz, these outliers were negligible, so participants were not discarded from the dataset.

Furthermore, we tested each participant’s visual acuity using Landolt C optotypes at 1.5 m distance. The average visual acuity in the sample was 1.62 (*SD* = 0.32, range = 1.2 to 2.6), reflecting sufficient vision. Since arithmetically averaged visual acuity values can be flawed, we applied a logarithmic averaging procedure to obtain the mean (Bach & Kommerell, 1998). That is, the logarithm of the individual visual acuity measures was obtained before averaging. Then, the antilogarithm of the subsequent average value was computed.

### Power analysis

Given a lack of suitable EEG studies, the sample size rationale emphasised maximising power for the behavioral effects of interest. To determine the smallest effect size of interest (Lakens, 2022) effect sizes from previous studies with similar research aims and experimental designs were considered. Our review revealed that effect sizes of earlier studies ranged between 0.16 to 0.84 *η*^2^_p_, the former effect being related to an interaction between probe congruency and memory conditions (single modality vs conjunction conditions) on response times (Daume et al., 2017)^1^. Thus, we determined our smallest effect size of interest to be 0.16 *η*^2^_p_. For an overview of the studies that informed this choice, please see the associated pre-registration at https://osf.io/q6wzr. Power analysis using MorePower 6.0 (Campbell & Thompson, 2012) targeting an effect size of 0.16 *η*2p, a power of 80%, and an alpha level of 0.05 for a 2 x 2 interaction resulted in a sample size estimate of 44 participants.

### Experimental setup and stimuli

The experiment was conducted in a dimly lit, sound-attenuated room (5.0 × 3.3 × 2.4 m³). The background noise level was kept below 20dB(A) using foam panels on the walls and ceiling and a woollen carpet on the floor. Stimulus presentation was governed using the E-Prime 3.0 software (Psychology Software Tools, Pittsburgh, PA). We used an AudioFile Stimulus Processor to control for the synchronisation of auditory stimuli with EEG triggers (Cambridge Research Systems, Rochester, UK). Additionally, we tested the synchronisation of auditory and visual stimuli. Visual onset timing was assessed using an analogue optical sensor, which was sensitive to the luminance change on the screen. To measure the onset of the auditory stimuli, the output of the AudioFile Stimulus Processor was fed into the EEG recording, using a NeurOne Isolation Box (Bittium Biosignals Ltd, Kuopio, Finland) for external analogue signals. This allowed us to record the sound signal transmitted to the loudspeakers and, thus, quantify the onset of the sound wave. According to our measurements, the two signals were highly synchronous with a negligible difference of ∼0.5 to 3 ms. Note that time measurements were obtained in a separate recording session, in which all combinations of visual and auditory stimuli were (dis)played.

Visual stimuli were displayed on a 49’’ centrally aligned 1800R curved monitor with a 5120 by 1440 pixel resolution and a 100 Hz refresh rate (Samsung, Seoul, South Korea). A full-range loudspeaker (SC 55.9 –8 Ohm; Visaton, Haan, Germany) was centrally mounted below the screen. Participants were seated in a comfortable chair at a distance of 130 cm from the screen.

As auditory stimuli, eight pure tones with frequencies ranging from 270 Hz to 3054 Hz (in steps of half an octave: 270, 381, 540, 763, 1080, 1527, 2159, 3054 Hz) were generated using the MATLAB® (R2022a) function ‘pure tone generator’ (Wojcicki, 2022). Tones had 20 ms fade-in and fade-out time windows with a sampling frequency of 44100 Hz and in full scale (amplitude = 1). A normalised auditory mask stimulus was generated by mixing all eight tones. As an auditory filler item, we created a white noise with an amplitude of 0.5 using free software called Audacity® (version 3.2.0). On average, auditory stimuli were presented at a sound level of 76 dB (LAeq, A-weighted, equivalent continuous sound level).

As visual stimuli, eight tear-drop-shaped orientations (RGB values 208, 208, 208) with angles between 22.5° to 337.5° (22.5°, 67.5°, 112.5°, 157.5°, 202.5°, 247.5°, 292.5°, 337.5°) were generated using the MATLAB function ‘polarplot’. The size of the tear-drop stimuli (from the tip to the opposite end) was 2°. A visual mask stimulus was generated by overlaying all eight orientations. Finally, as a visual filler item, we created a white blurred circle (RGB values 255, 255, 255 with a visual angle of 2.3°) using the MATLAB® (R2022a) function ‘polarplot’. All stimuli were presented in front of a light grey background (RGB values 128, 128, 128).

### Procedure, task, and experimental design

#### Audio-visual delayed match to sample task

Participants performed an audio-visual delayed match-to-sample task. The sequence of events within a trial is illustrated in Figure 1. A briefly flashing fixation cross (100 ms) signalled the start of each trial 1000 ms before the beginning of the trial sequence. Then, participants were presented with one or two sequentially displayed audio-visual memory items: spatially and temporally aligned tones, varying in frequency, and teardrops, varying in orientation. All memory items were presented in a central location (0° azimuth angle) for 200 ms. To minimise the impact of sensory memory traces, each memory item was followed by a centrally presented audio-visual mask for 200 ms. Memory items and masks were separated by an inter-stimulus interval (ISI) of 600 ms. Trials with one or two memory items were randomly intermixed. In one-item trials, an audio-visual filler stimulus replaced the first memory item to equate the overall trial length. Finally, following a delay period of 1200 ms, an audio-visual probe stimulus was displayed for 200 ms, and participants were asked to indicate whether the task-relevant probe features matched one of the items currently held in memory. Responses were recorded within a 2200 ms response period (relative to the onset of the probe) using a response pad with two horizontally arranged buttons. Each participant made button presses using left and right thumbs to respond. The assignment of response keys (left vs. right) to response options (‘yes’ vs. ‘no’) was counterbalanced across participants. Speed and accuracy were equally emphasised. Again, 2000 ms after probe offset, a brief flashing of the fixation cross (100 ms) signalled the start of the next trial.

**Figure 1.**
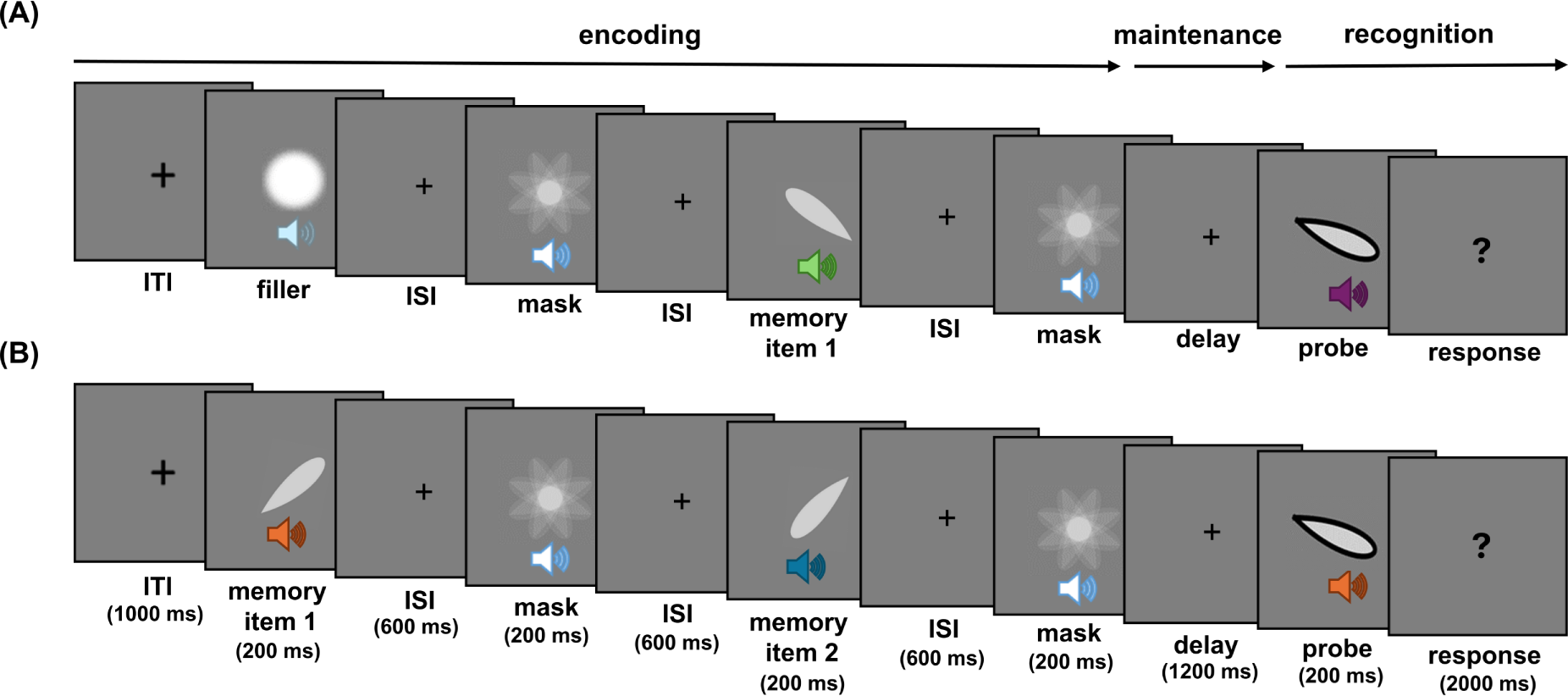
Schematic illustration of one versus two-item trials. *Note.* (A) In one-item trials, the first item was an audio-visual filler item; (B) while in two-item trials, this was replaced by a memory item. The three memory conditions only differed in terms of their task instructions, while the physical stimulation was always the same. *ITI = inter-trial interval, ISI = inter-stimulus interval*.

Overall, the experiment consisted of three blocked memory conditions: in the auditory condition, participants were instructed to attend only to the auditory features, while in the visual condition, they focused on only the visual features. In the following, the auditory and visual conditions are also referred to as the single-feature conditions. In the conjunction condition, participants were asked to attend to both the auditory and visual features and compare the whole object to the audio-visual probe features. Critically, participants were explicitly instructed in conjunction blocks to consider both features as part of the same object. It is important to note that the structure of the stimulus sequence remained identical in all three conditions, such that participants were always presented with audio-visual input. That is, the three memory conditions only differed in terms of their task instructions.

In addition to memory condition and set size, we also manipulated the type of probes. When combining stimulus features from two modalities, there are four resulting types of probes, depending on which features of the probes match or do not match the memory items. Essentially, those four probe types break down into two categories: (a) congruent probes, for which features in both modalities indicate a match or a no-match (i.e., auditory feature match + visual feature match; auditory feature no-match + visual feature no-match) and (b) incongruent probes, for which features in one modality indicate a match while the other modality indicates a no-match (i.e., auditory feature no-match + visual feature match; auditory feature match + visual feature no-match). Each probe type appeared equally often in the single-feature conditions (i.e., auditory and visual memory conditions) (see Figure 2A). This also results in an equal number of “yes” and “no” responses. However, considering both features were task-relevant in the conjunction condition, the proportion of trials per probe type had to be adjusted to balance the number of ‘yes’ and ‘no’ responses. Congruent-match trials (i.e., ‘auditory feature match + visual feature match’ probe) requiring a ‘yes’ response constituted 50% of trials in the conjunction condition. The other three probe types, which required a ‘no’ response in the conjunction condition, appeared equally often in 1/3rd each of the remaining 50% of trials (see Figure 2B). Across all memory conditions, the non-matching feature of the probe was always a novel stimulus, different from the features presented in each trial.

**Figure 2.**
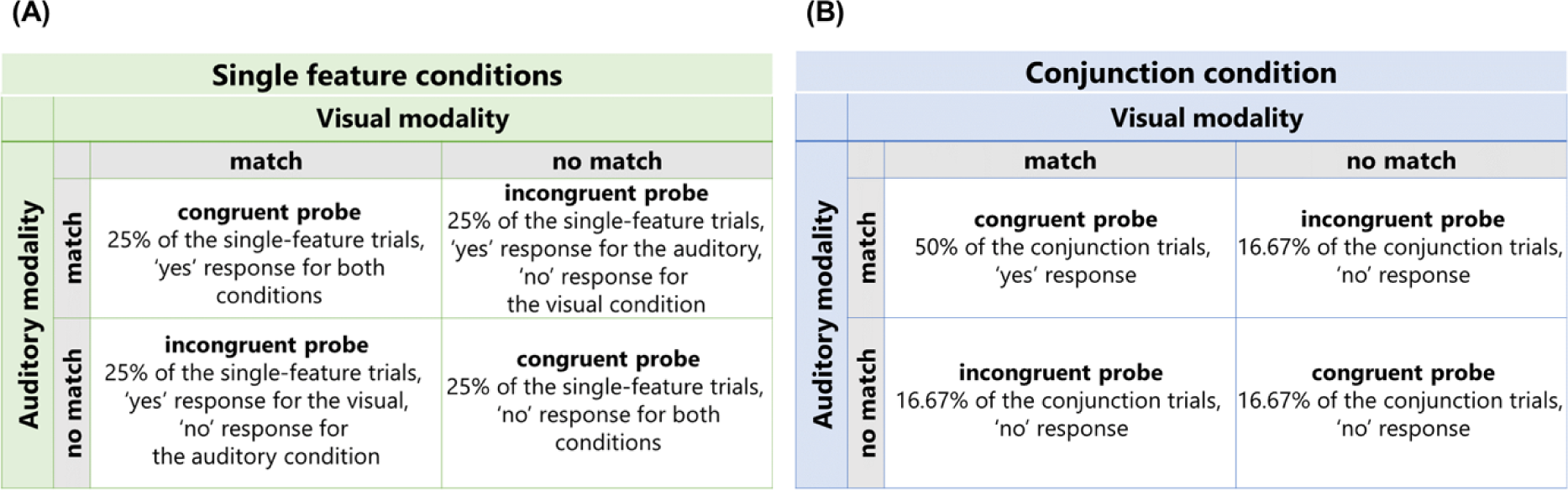
Illustration of probe types and respective proportion of trials per memory condition. *Note.* (A) In single-feature conditions, congruent and incongruent probe types appeared equally often; (B) to equalize ‘yes’ and ‘no’ responses across probe types in the conjunction condition, the ‘auditory match + visual match’ probe type was presented in half of the conjunction condition trials, while the other three probe conditions were equally distributed across the remaining 50% of trials.

Each memory condition included 240 trials (i.e., 120 one-item trials and 120 two-item trials), resulting in 720 trials. The experimental session was divided into 15 mini-blocks (five mini-blocks for each memory condition) of 48 trials. Each mini-block lasted approximately five minutes. The order of the mini-blocks was pseudorandomised such that memory conditions for each set of three blocks were randomly drawn from all three possible memory conditions until all conditions were chosen. The same procedure was followed for the next set of three mini-blocks, except that the first block of the next set could not have the same memory condition as the last block of the previous set had. Following each mini-block, participants were allowed to take a self-paced break. Preceding the start of the experimental session, participants practised the task by performing 24 trials for each memory condition. Overall, the entire task procedure (excluding EEG preparation time) took approximately one and a half hours.

#### Localizer task

A task-independent localizer procedure was employed to determine channels of interest for the time-frequency analysis. This procedure was primarily inspired by the localizer task conducted by (van Driel et al., 2014) and aimed at specifying electrodes that are maximally sensitive to auditory and visual stimuli processing. Notably, the localizer procedure enabled the selection of regions of interest independent of the experimental manipulations used in the main task. The procedure contained two blocks, an auditory and a visual block, each consisting of 80 trials. Participants were asked to attend to a series of randomly presented atonal tones in the auditory block. The tones were drawn from the same pool of eight sound stimuli presented during the main task, such that each sound was presented ten times. Likewise, in the visual block, participants attended to a series of randomly presented orientations drawn from the pool of stimuli presented during the main task, such that each exposure was repeated ten times. The auditory and visual stimuli were presented for 1000 ms with an inter-stimulus interval of 1000 ms. Participants were not required to respond to any of the stimuli presented. The procedure took approximately six minutes and always followed the main experiment. This task was only completed by a subset of participants (n = 24, 14 females, 2 participants left-handed).

### EEG recording

We used a 64 Ag/AgCl electrode cap (BrainCap; Brainvision, Gilching, Germany) with electrodes positioned according to the international 10-20 system (Pivik et al., 1993) to record EEG. The signal was sampled at 1000 Hz (NeurOne Tesla amplifier, Bittium Biosignals Ltd, Kuopio, Finland), and impedances were kept below twenty kΩ. The AFz and FCz electrodes served as ground and reference electrodes, respectively.

### EEG Preprocessing

#### Main experimental task

We used MATLAB® (R2022a) and the open-source toolbox EEGLAB (2022.1; Delorme & Makeig, 2004) for preprocessing. The continuous data were filtered offline using a 0.01 high-pass filter (filter order: 330001, transition bandwidth: 0.01 Hz, −6dB cutoff: 0.005 Hz) and a 40 Hz low-pass filter (filter order: 331, transition bandwidth: 10 Hz, −6dB cutoff: 45 Hz). Channels with a normalised kurtosis surpassing five standard deviations of the mean were rejected through an automated channel rejection procedure in EEGLAB (i.e., pop_rejchan function). On average, 4.64 channels were rejected per participant (range = 0 to 9, *SD* = 1.82). Removed channels were interpolated using the spherical spline of the neighbouring channels. Subsequently, the data were re-referenced to the average of all channels.

Next, we employed a semi-automatic artifact rejection method based on a rank-reduced independent component analysis (ICA). The data were down-sampled to 200 Hz to achieve a faster ICA decomposition. In addition, to increase the percentage of near-dipolar components, data were filtered with a 1 Hz high-pass filter (Winkler et al., 2015). Further, to reduce computation time, every other trial was selected after segmenting the data into epochs ranging from −1700 ms to 5500 ms relative to the onset of the first item. Finally, optimizing the signal-to-noise ratio in the ICA input, an automatic trial-rejection method was implemented (i.e., function pop_autorej) to exclude trials containing major artefacts and large voltage fluctuations (> 1000 μV). This procedure rejects trials, including data surpassing a given standard deviation in an iterative process (threshold: 5 SD, maximum percentage of total rejection per iteration: 5%).

The obtained ICA decomposition was then back-projected onto the original, filtered (0.01 Hz high-pass filter), and re-referenced continuous data with the initial sampling rate of 1 kHz. Again, the data were epoched from −1700 ms to 5500 ms relative to the onset of the first item and were baseline-corrected with a pre-stimulus baseline period of −700 to 0 ms. To identify artefactual independent components (ICs) reflecting eye movements, heart-or muscle activity, line noise, channel noise, or other non-brain activity, we used the EEGLab plugin ICLabel (v1.4; Pion-Tonachini et al., 2019). ICs that were given a probability estimate below 50% for the brain category were rejected from the dataset. Consequentially, 30.68 components (*SD* = 7.19) were excluded per participant on average. The remaining trials with large fluctuations (i.e., exceeding threshold limits of −150 to 150 μV) were excluded using the EEGLAB function ‘pop_eegthresh’. On average, 226.77 trials (*SD* = 17, 94.83%) remained in the auditory, 230.25 trials (SD = 16.07, 95.94%) in the visual, and 229.82 trials (*SD*= 16.87, 95.77%) in the conjunction condition.

#### Localizer task

We followed the abovementioned procedure in the same steps to pre-process the data from the localizer procedure. Given the shorter duration of each trial, we segmented the data into periods between −700 to 1500 ms relative to the onset of each experimental stimulus. Data were baseline-corrected with the pre-stimulus baseline period of −400 to −100 ms. On average, 3.19 channels were rejected per participant (range = 1 to 6, *SD* = 1.44), and 33.75 components (*SD* = 10.27, range = 16 to 58) were discarded. After pre-processing, on average, 78.29 trials (*SD* = 6.79, 97.86%) remained in the auditory, and 78.92 trials (*SD* = 5, 98.65%) in the visual condition, respectively.

### Time-frequency decomposition

To obtain event-related spectral perturbations (ERSPs; Makeig et al., 2004) and inter-trial phase clustering (ITPC, i.e., phase-locking factor; Tallon-Baudry et al., 1996) events, Morlet Wavelet Convolution was applied, using the EEGLAB routines calling the *pop_newtimef* function. Segmented EEG data from the main task and the localizer procedure were convolved with complex Morlet wavelets ranging in frequencies between 4 to 40 Hz (raising logarithmically in 52 steps). Aiming to obtain a good balance between temporal and frequency precision, the number of cycles increased by a factor of 0.5 with increasing frequency. At the lowest frequency, the number of cycles was 3, whereas at the highest frequency, it was 15.

For the main task data, the data were baseline-corrected, using a common trial average pre-stimulus baseline (dividing by average power across trials at each frequency, i.e., ‘gain model’), ranging from −600 to −200 ms before the onset of the first item. Compared to a condition-specific baseline correction, this procedure increases the signal-to-noise ratio in the baseline period. Before the common baseline procedure, a full-epoch length single-trial correction, was applied (Grandchamp & Delorme, 2011). The resulting ERSPs ranged from - 1282 to 5082 ms.

For the localizer task data, a condition-specific trial average pre-stimulus baseline was adopted, considering the entire pre-stimulus period (i.e., −282 to 0 ms) as a baseline. Again, the latter was preceded by a full-epoch length single-trial correction (Grandchamp & Delorme, 2011), divided by average power across trials at each frequency.

For the localizer task, ITPC was estimated to measure the consistency of phase angles over time across trials. As a result of the time-frequency decomposition described above, one phase angle value was obtained for each time-frequency point and trial. Those phase angles can be mathematically described as vectors with a unit length on a circle (Cohen, 2014). Averaging the phase vectors over trials, ITPC denotes the uniformity of the distribution of phase angles across trials at a given time point. ITPC can range from 0 to 1, reflecting no phase clustering over trials and perfect synchronization, respectively. The resulting ITPC time series ranged from −282 to 1082 ms relative to stimulus onset.

### Localizer-based channel selection

Regions of interest were defined by identifying those channels most strongly responding to the auditory and visual experimental stimuli, closely following the approach described by van Driel and colleagues (2014) for a similar purpose. To this end, inter-trial phase clustering activity in the theta (4-8 Hz) and alpha-band (8-12 Hz) are considered for the auditory and visual localizer task blocks, respectively (see Figure 3A).

**Figure 3.**
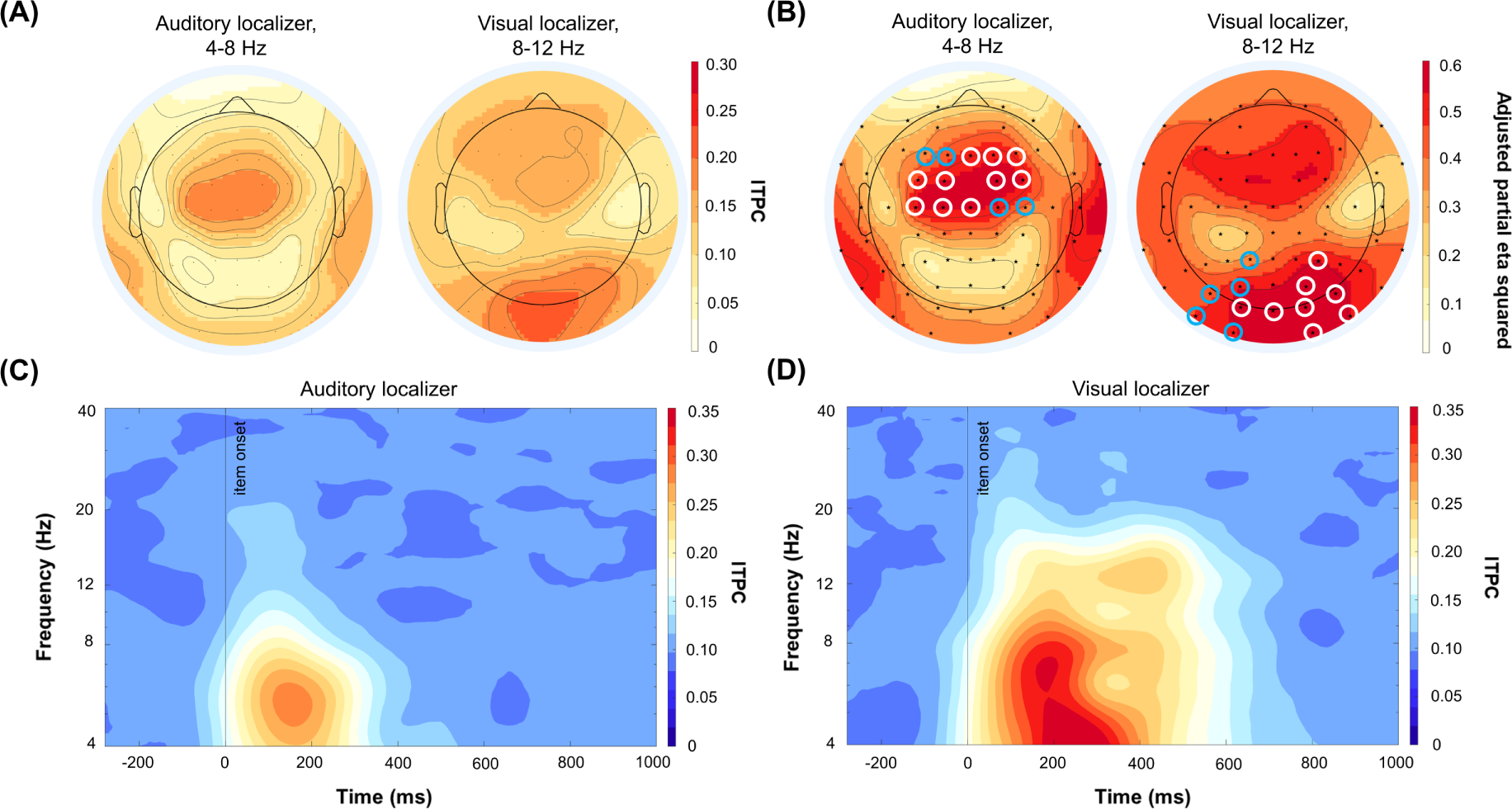
Task-independent localizer results and time-frequency dynamics of inter-trial phase clustering (ITPC) *Note.* (A) Localizer procedure showed a mid-central increase in theta (4-8 Hz) ITPC during the auditory stimuli and parieto-occipital increase in alpha (8-12 Hz) ITPC during visual stimuli presentation (101 to 384 ms post stimulus onset); (B) Electrodes of interest were selected based on the topographical distribution of effect sizes. Electrodes highlighted with white discs indicate channels exceeding the pre-defined threshold of 1 SD above the mean. Electrodes with blue discs were additionally selected to create symmetric electrode clusters; (C) Time-frequency dynamics of ITPC for the auditory stimuli show enhanced synchronization in the theta band; and (D) Time-frequency dynamics of ITPC for the visual stimuli show enhanced synchronization in the alpha-band.

To test for statistical differences in ITPC between baseline and activation periods, cluster-based permutation tests (Maris & Oostenveld, 2007), implemented in Fieldtrip (Version 20230422; Oostenveld et al., 2011) were performed. For a detailed description of the cluster-based permutation procedure, please see section ‘Cluster-based permutation tests’.

For the auditory condition, ITPC activity within the theta frequency range was contrasted between baseline (−282 to 0 ms relative to stimulus onset) and the post-stimulus activation period (101 to 384 ms relative to stimulus onset), as it was the dominant activity for this condition (see Figure 3C). In the visual condition, ITPC increases compared to baseline were evident in both the theta and the alpha-band (see Figure 3D). However, we only considered ITPC differences between baseline and activation within the alpha band. This choice was made since alpha activity is a particularly salient neural oscillation which is maximal in in the parieto-occipital regions (Zhigalov et al., 2019; Zhigalov & Jensen, 2020) and for comparability with the approach undertaken by van Driel and colleagues (2014). Data from all electrodes served as input to the cluster-based permutation test. Note that for both contrasts, the resulting significant clusters included all channels. Thus, the topographical distribution of effect sizes was obtained to allow for a meaningful selection of electrodes. Specifically, adjusted partial eta squared (*η*^2^_p_), as defined by Mordkoff (2019), was computed at each electrode. Finally, channels displaying an effect size greater than 1 standard deviation above the mean of effect sizes were selected. This resulted in a slightly right lateralized parieto-occipital cluster of channels for the visual condition (PO4/PO8/PO10, P4, O1/O2/O10/Oz) and one mid-central cluster of channels for the auditory condition (C1/C3/Cz, FC1/FC2, FC3/FC4, F2/F4/Fz). Considering that we did not have any a priori hypotheses about lateralization, we mirrored the channels around the anterior-posterior axis, yielding two channel clusters illustrated in Figure 3B.

### Statistical analyses

Statistical analyses were carried out using MATLAB® (R2022a) and JASP (version 0. 16.4.0). Unless otherwise specified, the significance of all tests was assessed at an alpha level of 0.05. For analyses of variance, we report Greenhouse-Geisser corrected *p*-values in case the assumption of sphericity was violated (Mauchley’s test *p* < .05). For repeated measures ANOVA and paired-sample t-tests, we report partial eta squared (*η*^2^_p_) and Cohen’s d as effect sizes, respectively. The standard deviation of the difference rates was used as a denominator to calculate Cohen’s d. Bonferroni-Holm method was used to correct multiple comparisons, and corrected p-values are described as *p_corr_*. Bayes Factors (BF_10_) were obtained as a supplementary measure, using JASP. According to Quintana and Williams (2018), a BF_10_ greater than 3, 10, or 30 is considered as moderate, strong, or very strong evidence in favor of the alternative hypothesis, respectively. A BF_10_ larger than 100 suggests extreme positive evidence. Analogously, a BF_10_ smaller than 0.33, 0.1, or 0.03 is considered as moderate, strong, or very strong evidence in favor of the null hypothesis, respectively, while a BF_10_ of smaller than 0.01 is indicative of extreme evidence for the null hypothesis. For cluster-based permutation tests (described in the section ‘Cluster-based permutation tests’), Cohen’s d is reported as the effect size measure. Here, we used the ‘average over cluster’ method to calculate effect sizes for significant clusters, averaging time-frequency data across the frequencies and time points that formed a cluster (Meyer et al., 2021). All cluster-based permutation tests were carried out using the MATLAB toolbox Fieldtrip (Version 20230422; Oostenveld et al., 2011; see section ‘Cluster-based permutation tests’).

#### Behavioral analyses

Mean reaction times (RTs) and mean accuracy (percentage of correct responses) were calculated to assess participants’ behavioural performance in the audio-visual delayed match-to-sample task. Responses that occurred after 2000 ms following the offset of the probe or occurred pre-maturely (i.e., within 150 ms following the onset of the probe) were excluded from further analyses.

##### Representational format: features vs. objects

First, for RTs and accuracy, a repeated-measures analysis of variance (rmANOVA) was performed, including the factors set size (set size 1 vs 2) and memory condition (auditory, visual, and conjunction conditions). This tackles the question to what extent an expected set size effect is driven by the number of features or the number of objects. In the single-feature conditions, the number of task-relevant features was one or two in set size 1 and set size 2 trials, respectively. In contrast, the number of task-relevant features in the conjunction condition was two or four in set size 1 and 2 trials, respectively (see Figure 4A). Thus, if only task-relevant information is stored in a purely feature-based fashion, the set size effect in the conjunction condition should be greater than in the single-feature conditions. In contrast, the number of objects is always one versus two in all three memory conditions (auditory, visual, and conjunction). Hence, a comparable set size effect across conditions would favour object-based storage. Notably, previous research has shown that working memory representations contain both a feature- and an object dimension (Li et al., 2022) hence, a less distinct pattern of results is always possible.

**Figure 4.**
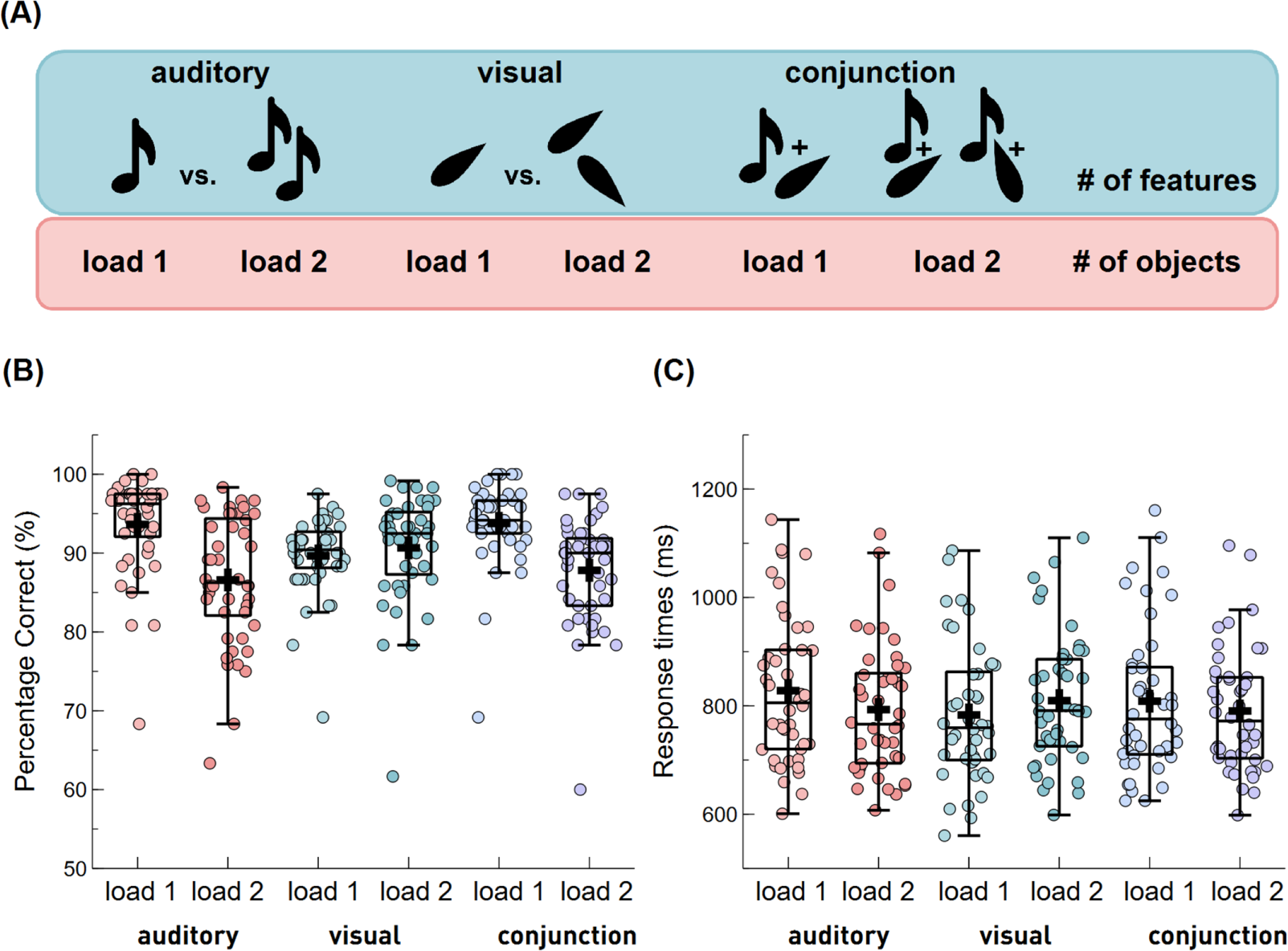
Set size effect across memory conditions. *Note.* (A) The upper row shows how many task-relevant_ features each set size (i.e., set size 1 vs set size 2) included per condition. The lower row illustrates how many objects each set size included per condition; (B) The proportion of correct responses varied by set size across memory conditions, (C) Reaction times varied by set-size across memory conditions. Boxplots show the +/-1.5 interquartile range and the median. The dots illustrate individual scores per condition. A black cross illustrates the condition mean.

##### Storage of task-irrelevant features

Second, to investigate to what extent the task-irrelevant feature of an attended object is still represented in working memory, we explored the effect of probe-congruency in the single-feature conditions. To this end, RT differences between trials with congruent probes (i.e., both probe-features match one of the memory items or both probe-features do not match one of the memory items) and incongruent probes (i.e., only one probe-features matches one of the memory items, whereas the other probe-feature has not been previously presented in a given trial) were computed. RT differences between congruent and incongruent probes served as the dependent variable for a 2 x 2 rmANOVA with set size (set size 1 vs 2) and condition (auditory vs visual) as within-subject factors. If the task-irrelevant feature is automatically encoded and subsequentially maintained in working memory in the auditory and the visual condition, despite the instruction to focus on input from only the task-relevant modality, responses should be faster for congruent compared to incongruent probes. Alternatively, suppose the information presented in the task-irrelevant modality is not encoded or filtered out after automatic encoding into working memory. In that case, the task-irrelevant probe feature dimension should not interfere with or affect performance. This may be modulated by memory load, such that under low working memory load (i.e., in one-item trials), both task-relevant and task-irrelevant features of a multisensory object are maintained. Thus, in one-item trials, we expected a stronger interference effect, reflected by lower accuracies and/or slower RTs in incongruent compared to congruent probe trials. In contrast, under a high working memory load (i.e., in two-item trials), we hypothesized that fewer working memory resources could be devoted to actively maintain features in the task-irrelevant modality, resulting in their more rapid decay or removal. If this is the case, congruent and incongruent probe trials should not have differed in RTs and accuracies due to weak representation of task-irrelevant features in two-item trials.

##### Task-relevancy of the interfering probe features

While the second analysis only focused on performance in the two single-feature conditions, an additional analysis contrasted the effect of probe congruency between the single-feature conditions and the conjunction condition. In the conjunction condition, any interference caused by an incongruent probe emerges from a task-relevant feature. In contrast, in the single-feature conditions, it is due to the interference from a task-irrelevant feature. Hence, this analysis allows us to test to what degree the probe congruency effect depends on the task relevance of the involved features. Critically, only trials requiring a ‘no’ response were included for these contrasts because ‘yes’ responses are typically faster than ‘no’ responses’ (Ratcliff & Hacker, 1981). This accounts for the fact that ‘yes’ and ‘no’ responses are not equally distributed across congruent and incongruent trials in the conjunction condition. A rmANOVA including the factors set size (set size 1 vs. 2) and memory condition (visual and conjunction conditions) was conducted to test for behavioral performance differences between the visual and conjunction conditions. The differences in RT and accuracy between congruent (visual feature no-match + auditory feature no-match) and incongruent (visual feature no-match + auditory feature match) probe trials served as the dependent variable, respectively. Another rmANOVA, including the factors set size (set size 2 vs 1) and memory condition (auditory and conjunction conditions), was conducted to contrast the auditory and the conjunction conditions. The differences in RT and accuracy between congruent (visual feature no-match + auditory feature no-match) and incongruent (auditory feature no-match + visual feature match) probe trials served as a dependent variable, respectively. Post hoc paired-sample t-tests were run to interpret significant interactions in the above analyses further. All comparisons were performed both for accuracy and reaction times data.

#### ERP analyses

The sustained anterior negativity (SAN) and the posterior negative slow wave (NSW) were chosen as EEG correlates of auditory and visual working memory load, respectively. Based on previous work, mean SAN amplitudes were calculated in a cluster of frontocentral electrodes, including AF3/4, F3/4, FC3/4, Fz, and FCz (Alunni-Menichini et al., 2014; Guimond et al., 2011). Mean NSW amplitudes were assessed in a cluster of parieto-occipital electrodes, including channels P7/8, P3/4, Pz, PO7/8, PO3/4, O1/2, and Oz (Diaz et al., 2021). Mean amplitudes were calculated during the maintenance period, ranging from 100 ms post-mask offset to probe onset (i.e., between 2700 ms to 3800 ms following the onset of the first item). The first 100 ms after mask offset were excluded to avoid including any sensory activity related to the sensory processing of the mask.

Mean amplitudes were submitted to a rmANOVA, including the within-subject factors memory conditions (auditory vs. visual vs. conjunction) and set size (set size 1 vs. 2). In addition, as outlined in the preregistration, to verify that the SAN and the NSW were sensitive to the set size manipulation of the task-relevant modality, one-sided paired sample t-tests were carried out, contrasting mean amplitudes between one- and two-item trials. Further, a series of pre-registered, planned contrasts were conducted: That is, paired-sample t-tests contrasting the set size effect (set size 2 vs. 1) between the memory conditions were performed (a) to verify whether SAN amplitudes were affected by the need to maintain concurrent visual features in the conjunction conditions and (b) to verify whether NSW amplitudes were modulated by the need to maintain concurrent auditory features in the conjunction condition. Finally, to observe if task-irrelevant features were actively maintained in working memory regardless of the task instructions, set size effects (i.e., paired-sample t-tests contrasting one- and two-item trials) were assessed for (a) SAN amplitudes in the visual condition and (b) NSW amplitudes in the auditory conditions.

#### Time-Frequency Analysis

Cluster-based permutation tests were conducted using the fieldtrip toolbox to identify frequency ranges and time windows sensitive to our experimental manipulations. Unless otherwise specified, default parameters were used (for details, see section ‘Cluster-based permutation tests’). As outlined before, to reduce the dimensionality of the data, we focused the analysis on two electrode clusters identified by a task-independent localizer procedure (for details, see section ‘Localizer-based channel selection’).

First, to test for the condition-dependent time-frequency modulations (irrespective of set size), the following pair-wise contrasts were created: visual vs. auditory, visual vs. conjunction and auditory vs. conjunction conditions. For both the anterior and posterior clusters, the time-frequency data was averaged across the two types of set size. Then, cluster-based permutation tests were run using all frequencies between 4 to 40 Hz and the entire epoch length, ranging from 1200 ms before and 5000 ms following the first item onset.

Second, the set size effect (set size 2 vs. 1) was tested separately for all three memory conditions. For both anterior and posterior clusters, time-frequency data were submitted to a series of cluster-based permutation tests, considering all frequencies (4 to 40 Hz) and the entire epoch duration (−1200 to 5000 ms).

Third, we were interested in the effect of probe congruency. Given that the behavioral analysis did not show any differences in the strength of the congruency effect between the single-feature conditions, the data were collapsed across the visual and the auditory condition. For both the anterior and posterior clusters, the time-frequency data were submitted to a cluster-based permutation test contrasting congruent versus incongruent probe trials. The statistical analysis included all frequencies (4 to 40 Hz) and all time points between the onset of the probe item and 1000 ms following the offset of the probe (3800 to 5000 ms). Earlier time points were not considered, because “congruency” is not known prior to the appearance of the probe. The conjunction condition was excluded from this analysis because both features were task-relevant. Finally, to test for an interaction between probe congruency and set size, analogous to the behavioural data analysis, the difference between congruent versus incongruent probe trials was computed separately for set size 1 and set size 2 trials and submitted to a cluster-based permutation test, including the same frequencies and time points as outlined above.

#### Cluster-based permutation tests

The fieldtrip-implemented cluster-based permutation test adheres to the following procedure: For each time-frequency pair (or time-frequency-channel sampling point for the localizer task data), a two-sided paired-sample t-test is conducted, contrasting the experimental conditions of interest. Only sampling points that yielded a *p*-value below .025 were considered for the subsequent clustering stage. At this stage, selected samples are clustered based on temporal, spectral (and spatial) adjacency. A triangulation method defined electrodes as neighbours for contrasts, including the electrode dimension. This method works by calculating a triangulation based on a two-dimensional projection of the channel positions and was used only for the localizer task data. The minimum number of neighbouring channels whose t-values were greater than the observed test statistic to form a cluster was three. The sum of all t-values serves as the cluster-level statistic for each identified cluster. If multiple clusters were identified, the maximum cluster-level statistic was selected for further evaluation of the statistical significance. Significance probabilities were calculated by using the Monte Carlo method. That is, all trials are randomly assigned a condition label (e.g., visual vs. auditory vs. conjunction conditions to test for the main effect of the condition). Following the random partitioning, a two-sided paired-sample t-test is conducted for each time-frequency pair (or time-frequency-channel sampling point). As for the observed data, the data are clustered based on temporal, spectral (and spatial) adjacency and a cluster-level statistic is computed. This random assignment of condition labels and the subsequent calculation of test statistics was repeated 1000 times, resulting in a time points x frequencies x participants x permutations matrix of cluster-test statistics. The observed test statistic is then compared to a threshold obtained from this distribution of test statistics under the null hypothesis. Differences between conditions were considered significant if the observed test statistic was lower than 1^st^ or larger than the 99^th^ percentile of the distribution of significance probabilities resulting from the permutation procedure. Please note that descriptions of the time range or frequency range comprised in a given cluster should not be interpreted as definite time or frequency boundaries of the true effect given that what is tested is the size of a cluster rather than exact time-frequency point differences between conditions (Sassenhagen & Draschkow, 2019).

## Results

### Behavioral results

#### Is the set size effect driven by the number of features or by the number of objects?

The first set of analyses considered condition (auditory vs. visual vs conjunction) as well as set size (set size 1 vs. set size 2) as independent variables, aiming at the question to what extent a potential set size effect (set size 1 – set size 2) is driven by the number of features or the number of objects. For RTs, rmANOVAs revealed no significant main effect of condition, *F*(2,1.44) = 1.69, *p_corr_* = .20, *η*^2^_p_ = 0.04, BF_10_ = 0.25, or set size, *F*(1, 43) = 1.26, *p* = .27, *η*^2^_p_ = 0.03, BF_10_ = 0.34. On the other hand, there was a greater accuracy for set size 1 compared to set size 2 trials, *F*(1, 43) = 58.74, *p* = < .001, *η*^2^_p_ = 0.58, BF_10_ > 1000. Similar to RTs, accuracies did not differ between conditions, *F*(2,1.32) = 0.61, *p_corr_* = .48, *η*^2^_p_ = 0.01, BF_10_ = 0.09. Critically, there was a significant interaction of condition and set size for RTs, *F*(2,1.59) = 16.21, *p_corr_* = < .001, *η*^2^_p_ = 0.27, BF_10_ = 23.97, as well as accuracies, *F*(2,1.35) = 43.58, *p*_corr_ = < .001, *η*^2^_p_ = 0.50, BF_10_ > 1000.

Follow-up paired-samples t-tests indicated that, compared to set size 2 trials, participants responded more accurately in set size 1 trials in the auditory*, t*(43) = 8.18, *p_corr_* =.003, *d* = 1.23, BF_10_ > 1000, and in the conjunction condition *t*(43) = 8.52, *p_corr_* = .002, *d* = 1.28, BF_10_ > 1000. In contrast, there was no significant set size effect for accuracy in the visual condition, *t*(43) = −1.50, *p_corr_* = .14, *d* = −0.23, BF_10_ = 0.46 (see Figure 4B).

Follow-up paired-samples *t*-tests showed that, compared to set size 1, set-size-2-trials in the visual condition resulted in slower RTs, *t*(43) = −2.57, *p_corr_* =.028, *d* = −0.39, BF_10_ = 2.98. In contrast, in the auditory condition, this pattern reversed, and participants responded faster in size-size-2 trials compared to set-size-1 trials, *t*(43) = 3.47, *p_corr_* =.003, *d* = 0.52, BF_10_ = 25.02. There was no significant effect of set size in RTs for the conjunction condition, *t*(43) = 1.77, *p_corr_* =.084, *d* = 0.27, BF_10_ = 0.68 (see Figure 4C).

Finally, a series of paired-samples t-tests were performed to see if the set size effect (set size 1 – set size 2) on both accuracy and RTs was significantly different between memory conditions. Firstly, pairwise comparisons between the auditory and the visual condition, showed that the set size effect in terms of RTs was greater for visual trials (*M* = −26.42, *SD* = 68.31) compared to auditory trials (*M* = 35.09, *SD* = 67.15), *t*(43) = 4.68, *p_corr_* =.003, *d* = 0.71, BF_10_ = 743.29. This is due to a reversed set size effect in the auditory condition (i.e., set size 2 trials were faster than set size 1 trials), while in the visual condition the set size effect was in the expected direction (i.e., set size 1 trials faster than set size 2 trials). For accuracies, we obtained a greater set size effect in the auditory (*M* = 7.05, *SD* = 5.72), compared to the visual condition (*M* = −1.02, *SD* = 4.51), *t*(43) = 6.97, *p_corr_* =.003, *d* = 1.05, BF_10_ > 1000, which was driven by greater accuracy in set size 1 trials compared to set size 2 trials in the auditory condition, while there was no significant set size effect in the visual condition (see above).

Secondly, for RTs, the set size effect (i.e., one-item faster than two-item trials) was greater in the visual (*M* = −26.42, *SD* = 68.31) compared to the conjunction condition (*M* = 18.06, S*D* =67.72), *t*(43) = −5.53, *p_corr_* =.002, *d* = −0.83, BF_10_ > 1000. On the contrary, the set size effect in terms of accuracy was greater in the conjunction conditions (*M* = 5.97, *SD* = 4.65) compared to the visual condition (*M* = −1.02, *SD* = 4.51), *t*(43) = −12.70, *p_corr_* =.002, *d* = −1.92, BF_10_ > 1000.

Finally, contrasting the set size effect between the auditory and the conjunction condition, we did not find any significance differences for RTs, *t*(43) = 1.46, *p_corr_* =.15, *d* = 0.22, BF_10_ = 0.44, or accuracies, *t*(43) = 1.08, *p_corr_* =.29, *d* = 0.16, BF_10_ = 0.28.

With respect to the question of whether audio-visual object storage is feature- or object based, the data shows no perfectly consistent pattern. While auditory vs. conjunction contrast is in line with an object-based account of working memory, the visual vs. conjunction contrast, at least in terms of accuracy, rather favors a feature-based account.

#### Is the task-irrelevant feature of an attended object stored in working memory?

The second set of analyses aimed at assessing whether task-irrelevant cross-modal features were represented in working memory, thus interfering with performance at recall. As mentioned before, the conjunction condition was excluded from these analyses, given that both probe dimensions were task-relevant. RTs and accuracy differences between congruent and incongruent probe trials served as the dependent variables for two separate rmANOVA including the factors condition (auditory, visual) and set size (set size 1 vs. 2). There was a main effect of set size for RTs, *F*(1, 43) = 7.83, *p* = .008, *η*^2^_p_ = 0.15, BF_10_ =5.08, pointing out greater RTs differences between congruent vs incongruent trial types in set size 1 compared to set size 2 trials. Additional paired-samples t-tests revealed that, compared to incongruent trials, participants performed faster in congruent probe trials in set size 1, *t*(43) = −3.05, *p_corr_* =.008, *d* = −0.46, BF_10_ = 8.96. This congruency effect vanished in set size 2 trials, *t*(43) = 0.93, *p_corr_* = .36, *d* = 0.14, BF_10_ = 0.25. Neither the main effect of condition, *F*(1, 43) = 0.17, *p* = .69, *η*^2^_p_ = 0.004, BF = 0.21, nor the interaction of condition and set size, *F*(1, 43) = 0.009, *p* = .93, *η*^2^_p_ = 0.002, BF = 0.70, were significant (see Figure 5A).

**Figure 5.**
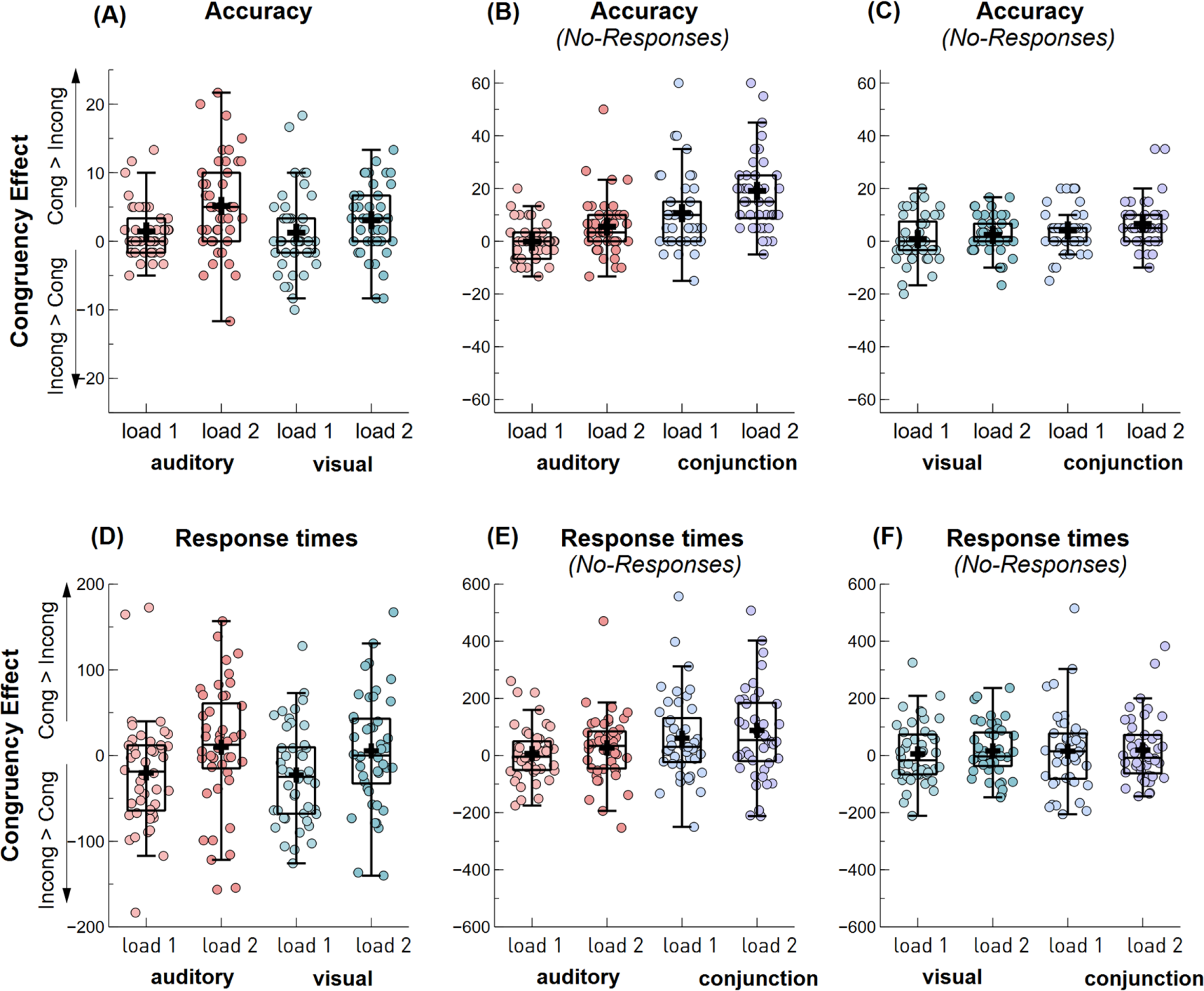
Probe congruency effect between memory conditions for different set trials. *Note.* (A) Proportion of correct responses for the congruency effect (congruent – incongruent) between auditory and visual conditions; (B) Proportion of correct responses for the congruency effect between auditory and conjunction conditions, collapsed over trials requiring a ‘no’ response. The data depicted for the conjunction condition refers to trials with a probe containing an auditory-no-match feature paired with a visual-match feature; (C) Proportion of correct responses for the congruency effect between visual and conjunction conditions, averaged over trials requiring a ‘no’ response. The data depicted for the conjunction condition refers to trials with a probe containing a visual-no-match feature paired with an auditory-match feature; (D) RTs for the congruency effect between auditory and visual conditions; (E) RTs for the congruency effect between auditory and conjunction conditions, collapsed over trials requiring a ‘no’ response. The data depicted for the conjunction condition refers to trials with a probe containing an auditory-no-match feature paired with a visual-match feature; (F) RTs for the congruency effect between visual and conjunction conditions, averaged over trials requiring a ‘no’ response. The data depicted for the conjunction condition refers to trials with a probe containing a visual-no-match feature paired with an auditory-match feature. Boxplots show the +/- 1.5 interquartile range and the median. The dots illustrate individual participant averages per condition. A black cross illustrates the condition mean.

For accuracy, a significant main effect of set size, *F*(1, 43) = 13.39, *p* = .008, *η*^2^_p_ = 0.24, BF_10_ = 19.71, pointed out greater accuracy differences between congruent vs. incongruent trial types in set size 2, compared to set size 1 trials. Additional paired-samples t-tests revealed that, compared to incongruent trials, participants scored higher in congruent probe trials for both set size 1, *t*(43) = 2.41, *p_corr_* = .02, *d* = 0.36, BF_10_ = 2.16, and set size 2, *t*(43) = 6.11, *p_corr_* = .002, *d* = 0.92, BF_10_ > 1000. Further, there was neither a significant interaction of condition and set size, *F*(1, 43) = 1.47, *p* = .23, *η*^2^_p_ = 0.033, BF_10_ = 0.56, nor a main effect of condition, *F*(1, 43) = 1.91, *p* = .17, *η*^2^_p_ = 0.04, BF_10_ = 0.42 (see Figure 5D). Overall, these results show that incongruent probes decreased accuracy in both set size conditions and increased RTs in one-item-trials. This suggests that task-irrelevant features are stored in working memory to some degree, regardless of the task goals.

#### Is the probe congruency effect affected by the task-relevance of a mis-matching feature?

Finally, to explore whether the probe congruency effect is modulated by the task relevance of the interference-inducing feature of the probe, separate sets of rmANOVAs were conducted, contrasting the conjunction condition with the single-feature conditions. As outlined in section ‘Task-relevancy of the interfering probe features’, only trials requiring a no-response were included in this analysis. First, two rmANOVA were performed, including the factors memory condition (visual vs. conjunction) and set size (1 vs. 2) on the RT and accuracy differences between congruent (visual feature no-match + auditory feature no-match) and incongruent (visual feature no-match + auditory feature match) probe trials. The RT analysis yielded no significant main effects, nor an interaction (all *p* > .68) (see Figure 5F). Yet, accuracies for memory conditions significantly differed from each other, *F*(1, 43) = 6.82, *p* = .012, *η*^2^_p_ = 0.06, BF_10_ = 3.72, indicating a greater probe congruency effect for the conjunction condition (*M* = 5.34, *SD* = 6.64), compared to the visual condition (*M* = 1.63, *SD* = 5.98) (see Figure 5C). This result suggests that greater interference results from a task-relevant but non-matching feature, leading to lower accuracy. Planned follow-up contrasts verified the presence of an interference effect (i.e., congruency effect) in the conjunction condition, *t*(43) = 5.34, *p* < .001, *d* = 0.81, BF_10_ > 1000, but not in the visual condition, *t*(43) = 1.81, *p* = .08, *d* = 0.27, BF_10_ = 0.73. Neither the main effect of set size, *F*(1, 43) = 2.53, *p* =.12, *η*^2^_p_ = 0.06, BF_10_ = 0.59, nor the interaction of condition and set size in accuracies, *F*(1, 43) = 0.18, *p* =.68, *η*^2^_p_ = 0.004, BF_10_ = 0.24, was significant.

A second group of rmANOVA was run with factors memory conditions (auditory vs. conjunction) and set size (1 vs. 2). Again, the RT and accuracy differences between congruent (auditory feature no-match + visual feature no-match) vs. incongruent (auditory feature no-match + visual feature match) probe trials served as dependent variables. A significant main effect of condition in RTs, *F*(1, 43) = 7.21, *p* =.010, *η*^2^_p_ = 0.07, BF_10_ = 4.56, indicates a greater probe congruency effect for the conjunction condition (*M* = 74.21, *SD* = 138.83) compared to the auditory condition (*M* = 16.90, *SD* = 82.97), reflecting slower RTs in the congruent compared to incongruent trials (see Figure 5E). Planned follow-up contrasts established the congruency effect in the conjunction condition, *t*(43) = 3.55, *p* < .001, *d* = 0.54, BF_10_ = 30.78, but not in the auditory condition, *t*(43) = 1.35, *p* = .18, *d* = 0.20, BF_10_ = 0.38. Further, there was neither a significant interaction of condition and set size, *F*(1, 43) = 0.06, *p* = .81, *η*^2^_p_ = 0.25, BF_10_ = 0.23, nor a main effect of set size, *F*(1, 43) = 1.53, *p* = .22, *η*^2^_p_ = 0.01, BF_10_ = 0.40.

Similar to the RT differences between the two conditions, a significant main effect of memory condition on the accuracy, *F*(1, 43) = 36.51, *p* < .001, *η*^2^_p_ = 0.28, BF_10_ > 1000, showed a greater probe congruency effect for the conjunction condition (*M* = 14.89, *SD* = 13.83) compared to the auditory condition (*M* = 2.69, *SD* = 7.44), reflecting impaired performance for incongruent compared to congruent trials (see Figure 5B). Planned contrasts verified the congruency effect both in the auditory, *t*(43) = 2.40, *p* = .02, *d* = 0.36, BF_10_ = 2.11, and in the conjunction conditions, *t*(43) = 7.14, *p* < .001, *d* = 1.08, BF_10_ >1000. Additionally, there was a significant main effect of the set size, *F*(1, 43) = 24.26, *p* < .001, *η*^2^_p_ = 0.09, BF_10_ = 578.94, indicating a stronger probe congruency effect for a higher memory load. Yet, the interaction of the two factors was not significant, *F*(1, 43) = 1.05, *p* = .31, *η*^2^_p_ = 0.003, BF_10_ = 0.33. Overall, accuracy results suggest that the interference from the non-matching feature of the attended object exists in case of this feature is task relevant rather than task irrelevant.

## ERP data

ERP correlates of working memory storage were investigated to track the maintenance of task-relevant and task-irrelevant features (see Figure 6A for the respective electrodes used in the analyses). rmANOVAs, including the factors memory condition (auditory vs. visual vs. conjunction) and set size (1 vs. 2), were conducted for both SAN (auditory working memory storage) and NSW (visual working memory storage) amplitudes, respectively.

**Figure 6.**
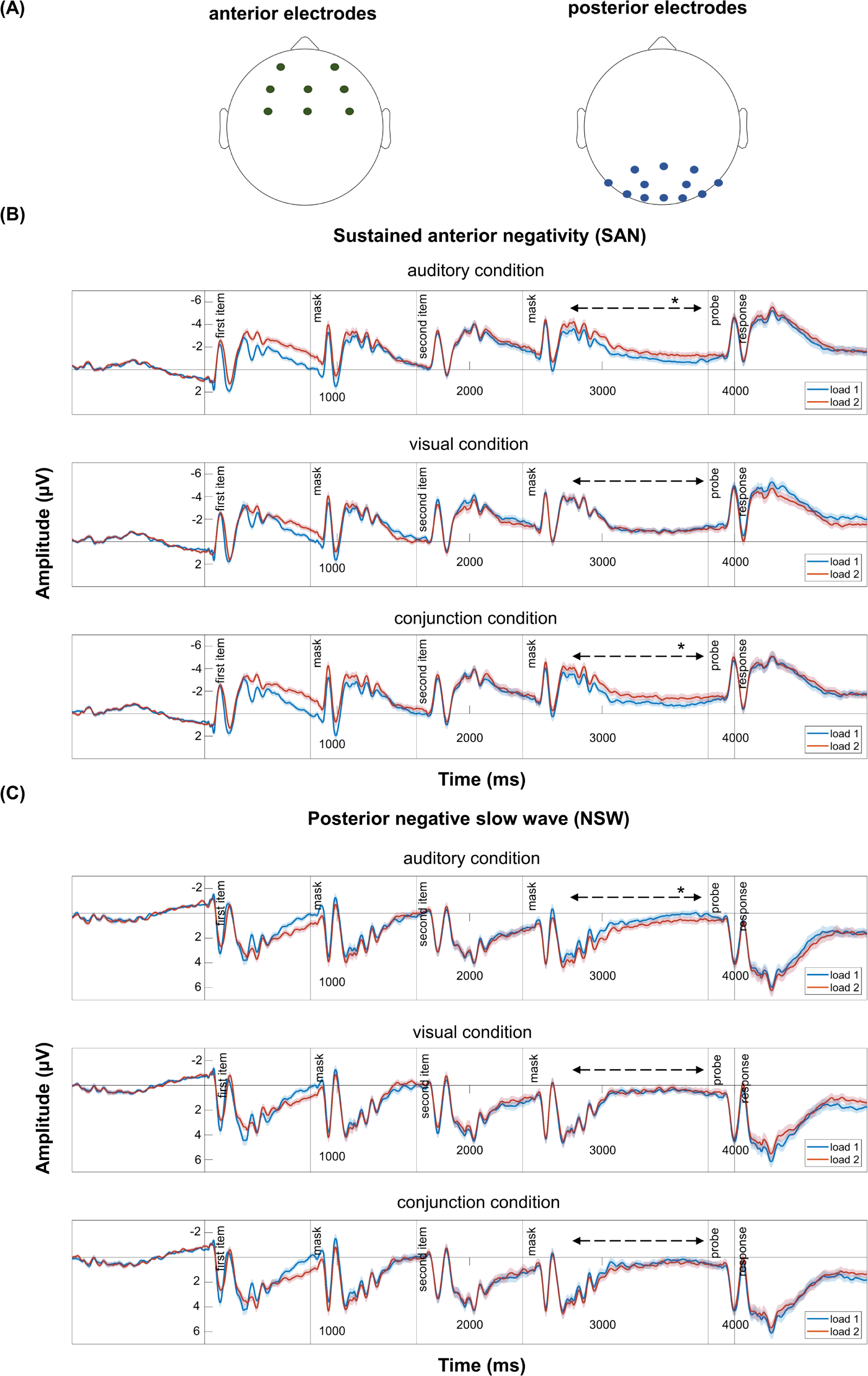
Condition-specific, grand average ERP waveforms at anterior (SAN) and posterior (NSW) electrode sites. *Note.* (A) The topographies show the electrode clusters of interest, left panel for the anterior clusters used for SAN and right panel for the electrodes used for NSW. The ERPs in (B) and (C) depict the time course of the SAN and NSW, respectively, separately for the auditory, visual, and conjunction condition and for set size 1 (blue lines) and set size 2 trials (red lines). Black arrows indicate the corresponding analysis time window, spanning the maintenance period. Significant paired-sample t-test contrasts (i.e., auditory set size 1 vs. set size 2 and conjunction set size 1 vs. set size 2 comparisons for SAN and auditory set size 1 vs. set size 2 for NSW) were marked with an asterisk (*p* < 0.05).

An analysis of mean SAN amplitudes revealed a main effect of set size for SAN, *F*(1, 43) = 7.37, *p* = .010, *η*^2^_p_ = 0.15, BF_10_ = 2.91. There was no significant main effect of condition, *F*(1, 43) = 0.71, *p* = .49, *η*^2^_p_ = 0.02, BF_10_ = 0.08, nor a significant interaction, *F*(1, 43) = 2.54, *p* = .09, *η*^2^_p_ = 0.06, BF_10_ = 1.16. Despite the lack of a significant interaction, pre-registered one-sided paired-samples t-tests were conducted to verify the presence of a set size effect within conditions (see Figure 6B). Accordingly, set size 2 trials displayed significantly more negative SAN amplitudes than set size 1 trials in both the auditory, *t(*43) = 2.82, *p* = .007, *d* = 0.43, BF_10_ = 5.21, and conjunction conditions, *t*(43) = 2.26, *p* = .03, *d* = 0.34, BF_10_ = 1.60, in which auditory features were task-relevant. However, no effect of set size was evident in the visual condition, *t*(43) = −0.12, *p* = .91, *d* = −0.02, BF_10_ = 0.16, where auditory features were task-irrelevant. The corresponding Bayes Factor provides moderate evidence for the null hypothesis, indicating that task-irrelevant features are not actively maintained.

An analysis of mean NSW amplitudes revealed no significant effects (all *p* > .05). Again, despite the lack of a significant interaction, planned one-sided paired-sample t-tests were conducted as pre-registered (see Figure 6C). Contrary to our expectations, no significant set size variation of NSW amplitudes was evident in the visual, *t*(43) = 0.25, *p* = .81, *d* = 0.04, BF_10_ = 0.17, or the conjunction, *t*(43) = −0.98, *p* = .33, *d* = −0.15, BF_10_ = 0.26, condition, where visual features where task-relevant. In the auditory condition, NSW amplitudes were significantly more positive in set size 2 trials than in set size 1 trials, *t*(43) = −2.65, *p* = .01, *d* = −0.4, BF_10_ =3.53, exhibiting a reversed set size effect.

### Time-Frequency results

#### The effect of memory condition

Figure 7 shows the time course of oscillatory power for all three memory conditions for anterior (see Figure 7A) and posterior clusters (see Figure 7C), collapsed across trials with different set sizes, and the respective difference plots (see Figure 7B, 7D). Inspecting the time course of oscillatory power, a similar pattern emerges across all three memory conditions: In response to the item and mask presentations, a stimulus-evoked increase in the theta band and a decreased alpha and beta band power is evident. Shortly after encoding and in the delay interval, this activity was followed by an increase in power in the upper alpha and lower beta band. After the probe presentation, a prominent decrease in alpha- and lower beta power and an increase in theta power can be observed (see Figure 7A, 7C).

**Figure 7.**
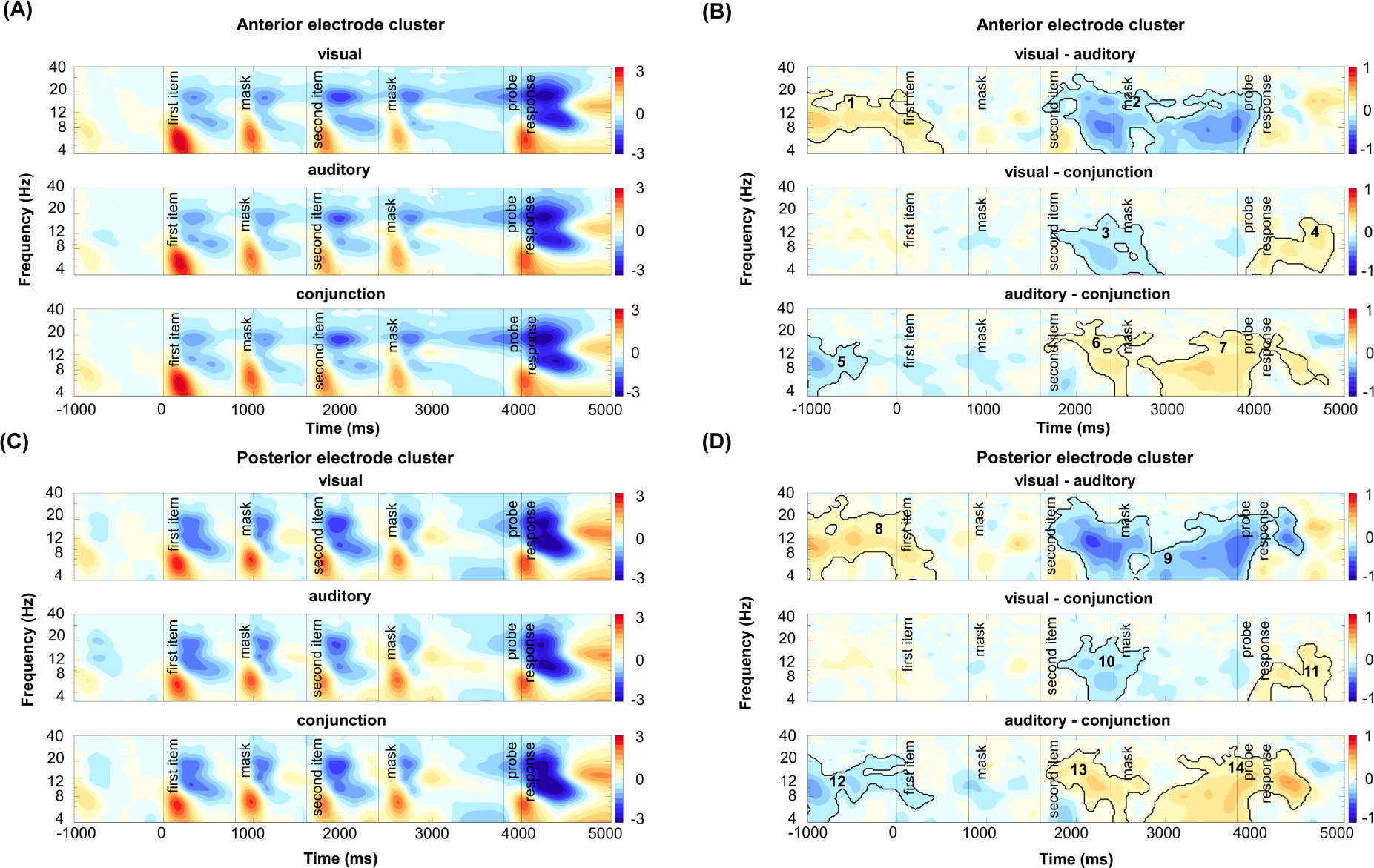
Time-frequency modulations across memory conditions and cluster-based permutation test results. *Note.* (A) Power was averaged across the midcentral electrode cluster including C1/C3/Cz, FC1/FC2, FC3/FC4, F2/F4/Fz, FT10 for each condition; (B) Differences between conditions were tested with cluster-based permutation tests; (C) Power was averaged across the parieto-occipital electrode cluster including PO4/PO8/PO10, P4, O1/O2/O10/Oz for each condition; (D) Differences between conditions were tested with cluster-based permutation tests. Solid lines illustrate significant clusters with a t-mass value smaller than 1st or larger than the 99th percentile of the distribution of significance probabilities of the null distribution.

Cluster-based permutation analyses, contrasting the pair-wise combinations of memory conditions, revealed significant differences between conditions in the encoding period after the second-item presentation, in the maintenance interval, as well as during probe presentations. Critically, the pattern of results was highly comparable for the anterior and the posterior clusters (see Figure 7B, 7D).

In the encoding period following the second-item presentation, differences between the conditions appear most prominent in the alpha band but expand into the adjacent theta and beta bands. Specifically, the contrast between the visual and the conjunction condition shows a relative increase in power in the conjunction condition following the second memory item. The corresponding cluster (cluster 3 in anterior electrodes: *p* = .001, *d* = −0.17 and cluster 10 in posterior electrodes: *p* = .001, *d* = −0.17; see Figure 7B and 7D) comprises the theta- and alpha range while also extending into the lower beta-band (anterior electrodes: 4 to 19 Hz, posterior electrodes: 4 to 22 Hz). In the time domain, the cluster in the observed data extends from 1740 to 2970 ms (relative to first item-onset) at anterior electrodes and from 1800 to 2800 ms at posterior electrodes. In the response period, the cluster-based permutation analysis also revealed a relatively stronger power suppression in the conjunction condition compared to the visual condition. The corresponding cluster (cluster 4 in anterior electrodes: *p* = .001, *d* = 0.18, cluster 11 in posterior electrodes: *p* = .001, *d* = 0.18; see Figure 7B and 7D) mainly comprises the theta- and alpha-band (anterior electrodes: 4 to 18 Hz, posterior electrodes: 4 to 18 Hz) and extends roughly from 3910 to 4870 ms at anterior and from 3930 to 4850 ms at posterior electrodes.

When comparing the auditory and the conjunction conditions, a relative increase in power can be observed in the auditory compared to the conjunction condition in the encoding period following the second-item presentation. The corresponding cluster in the observed data (cluster 6 in anterior electrodes: *p* = .001, *d* = 0.17, cluster 13 in posterior electrodes: *p* = .003, *d* = 0.21; see Figure 7B and 7D) comprises the alpha-band but extends into the theta and lower beta-band (anterior electrodes: 4 to 29 Hz, posterior electrodes: 4 to 27 Hz). A second cluster (cluster 7 in anterior electrodes: *p* = .001, *d* = 0.22, cluster 14 in posterior electrodes: *p* = .001, *d* = 0.26; see Figure 7B and 7D) emerges in the maintenance period and extends into the recall period (anterior electrodes: 2700 to 4800 ms, posterior electrodes 2760 to 4610 ms). Again, the cluster spans a broad range of frequencies in the theta, alpha, and lower beta-band (anterior electrodes: 4 to 22 Hz, posterior electrodes: 4 to 29 Hz). The latter cluster indicates a relative increase in power in the auditory compared to the conjunction condition, or in other words, a relatively stronger decrease in power in the conjunction condition.

Finally, when contrasting the auditory and the visual conditions, the cluster test reveals a broad cluster (cluster 2 in anterior electrodes: *p* = .001, *d* = −0.26; cluster 9 in posterior electrodes: *p* = .001, *d* = −0.30; see Figure 7B and 7D) that spans the encoding interval after second-item presentation, the maintenance period, and the response interval (anterior electrodes: 1630 to 4060 ms, posterior electrodes 1700 to 4530 ms). In the frequency dimension, the effect appears most prominent in the alpha-band but clearly extends broadly in the adjacent theta and beta band (anterior electrodes: 4 to 32 Hz, posterior electrodes: 4 to 35 Hz). This cluster reflects a sustained relative increase in power in the auditory condition compared to the visual condition.

Lastly, the cluster tests also revealed significant differences in the baseline period, which we mainly attributed to the block-wise presentation of conditions. Namely, there was a stronger anticipatory suppression of alpha power in the auditory condition compared to the conjunction and visual conditions, respectively. In the former case, the corresponding cluster (cluster 5 in anterior electrodes: *p* = .001, *d* = - 0.19, cluster 12 in posterior electrodes: *p* = .001, *d* = −0.20; see Figure 7B and 7D) comprised time points from −1280 to −350 ms at anterior electrodes and from −1280 to 400 ms at posterior electrodes. In the frequency domain, the cluster spanned predominantly the alpha band but expanded into the adjacent theta and lower beta-band (anterior electrodes: 4 to 30 Hz, posterior electrodes: 4 to 28 Hz). In the latter case, the corresponding cluster (cluster 1 in anterior electrodes, *p* = .001, *d* = 0.21, cluster 8 in posterior electrodes: *p* = .001, *d* = 0.22; see Figure 7B and 7D) comprised time points from - 1280 to 505 ms at anterior electrodes and from −1280 to 420 ms at posterior electrodes. In the frequency domain, the effect was most prominent in the alpha band, extending through theta and beta bands (anterior electrodes: 4 to 22 Hz, posterior electrodes: 4 to 36 Hz). Notably, the clusters in the post-stimulus period cannot be due to those baseline differences, as the direction of effects in the baseline is reversed compared to those found later in the trial.

In summary, the three pair-wise contrasts outlined above show a prominent relative increase in alpha power (and adjacent frequency bands) in auditory blocks compared to visual and conjunction blocks in the encoding and maintenance period. Further, a relative increase in alpha power (as well as adjacent frequency bands) was evident in the conjunction compared to visual blocks, during maintenance. Finally, a consistent decrease in alpha power (and adjacent frequency bands) in the conjunction condition was observed at recall, compared to both single-feature conditions.

#### The effect of memory load

Figure 8 shows the pair-wise combinations of set size 2 vs. 1 trial within each memory condition, for the anterior (see Figure 8A) and the posterior electrode cluster (see Figure 8B). Cluster-based permutation analyses showed significant differences at early stages following the first item (i.e., filler item in set size 1 trials and first memory item in set size 2 trials), the mask, and the second item (i.e., first memory item in set size 1 trials and second memory item in set size 2 trials) together with significant differences at recall. Although the specific time-frequency boundaries of the obtained clusters differed slightly for the different memory conditions and electrode clusters, we can differentiate three major sets of functionally compatible clusters:

(i) In the first few hundred milliseconds of the trial, set size 2 trials elicit a stronger decrease in theta power compared to set size 1 trials. This effect is only significant at the anterior electrode cluster (visual: *p* = .001, *d* = −0.42, auditory: *p* = .002, *d* = −0.49, conjunction: *p* = .007, *d* = −0.51), and extends to later time points (∼900 ms) and higher frequencies (∼4-28 Hz) in the visual condition. This set of clusters (see cluster 2,6,8, Figure 8A) can be attributed to differences in processing between the filler item in set size 1 trials and the encoding of a memory item in set size 2 trials and is thus not of major interest here.
(ii) Following the presentation of the first mask stimulus, a relatively sustained (∼700 to 1600 ms) broad-band power increase spanning the theta, alpha, and lower beta-band is evident in set size 2 compared to set size 1 trials. This effect (see clusters 11, 13,15 in Figure 8B) is predominantly reflected at posterior electrode sites in the visual (*p* = .001, *d* = 0.27), auditory (*p* = .001, *d* = 0.29), and conjunction condition (*p* = .002, *d* = 0.28; but see also cluster 7 in figure 8A for a slightly earlier, but likely related cluster in the auditory condition at anterior electrode sites). This set of effects is followed by a relative increase in theta power in set size 2 compared to set size 1 trials. Corresponding, significant clusters are in the visual condition (*p* = .001, *d* = −0.28) and in the conjunction condition (*p* = .001, *d* = 0.36) at anterior electrode sites (cluster 3 and 9 in Figure 8A). These two sets of clusters can – again – be attributed to differences in required cognitive control and attentional processing when processing the filler item in set size 1 trials compared to the storage and maintenance of the first memory item in set size 2 trials. Again, these effects were not a main objective of this investigation.
(iii) Finally, at recall, set size 2 trials consistently result in a stronger decrease in alpha power (and adjacent frequencies) compared to set size 1 trials. Corresponding, significant clusters were identified in all memory conditions and at both electrode sites (anterior electrodes sites: visual condition: *p* = .001, *d* = −0.28, conjunction condition: *p* = .001, *d* = −0.31; posterior electrode sites: visual condition: *p* = .001, *d* = −0.35, auditory condition: *p* = .002, *d* = −0.29, conjunction condition: *p* = .001, *d* = −0.33), except for the auditory condition at the anterior electrode site (*p* = .013). This set of clusters (see clusters 4, 10, 12, 14, 16 in Figure 8A, 8B) can be attributed to greater attentional demands when retrieving two-item working memory representations compared to one-item working memory representations.

**Figure 8.**
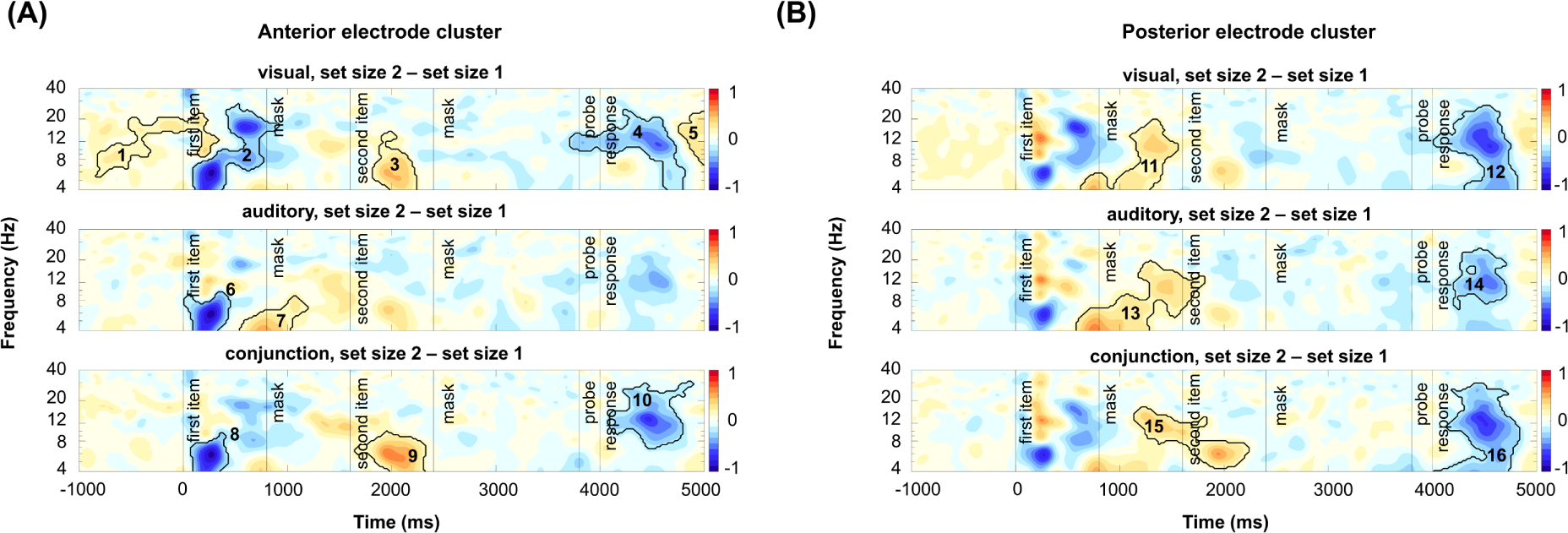
Cluster-based permutation test results for set size 2 vs. 1 contrast across memory conditions. *Note.* (A) Pair-wise contrasts (between set size 2 vs set size 1 trials) for time-frequency power between conditions were tested with cluster-based permutation tests for the mid-central electrodes; (B) Pair-wise contrasts (between set size 2 vs set size 1 trials) for time-frequency power between conditions were tested with cluster-based permutation tests for the parieto-occipital electrodes. Solid lines illustrate significant clusters with a t-mass value smaller than 1st or larger than the 99th percentile of the distribution of significance probabilities of the null distribution.

Unlike other contrasts, a significant cluster in the baseline period of the visual condition was observed at anterior electrodes (cluster 1 in Figure 8A). This effect was not further interpreted given that it was not possible to prepare for a two-item trial until the first item appeared in a given trial.

In summary, differences at early stages point to the processing of the filler item in one-item trials compared to the encoding of the first memory item in two-item trials. Critically, the relative power increase in theta range might point out filler-item processing, which is later discarded, as addressed by a relative decrease in theta range. Furthermore, stronger alpha power suppression at recall addresses greater attentional demands when retrieving two-item working memory representations compared to one-item working memory representations.

#### The effect of probe congruency

Furthermore, we were interested in the effect of probe congruency. This was an exploratory analysis as a follow-up on the interaction between these factors in behavioral performance. Since the behavioral analysis did not show an interaction of probe congruency and condition, the data were average across the two single-feature conditions. Figure 9 shows a probe congruency effect on time-frequency modulations, indicated by a stronger alpha power suppression following incongruent probes, compared to congruent probes. The corresponding cluster in the observed data (cluster 1 in anterior electrodes: *p* = .001, *d* = −0.19, cluster 2 in posterior electrodes: *p* = .001, *d* = −0.19) comprises the alpha-band but extends into the theta and lower beta-band (anterior electrodes: 7 to 24 Hz, posterior electrodes: 4 to 24 Hz), sustained between 4290 to 5000 ms at anterior and 4290 to 5000 ms at posterior electrodes. This result suggests a higher need for attentional control for incongruent probes, also supported by behavioral performance decay. The analysis did not show any significant interaction between set size and probe congruency (all *p* > .22, data not shown).

**Figure 9.**
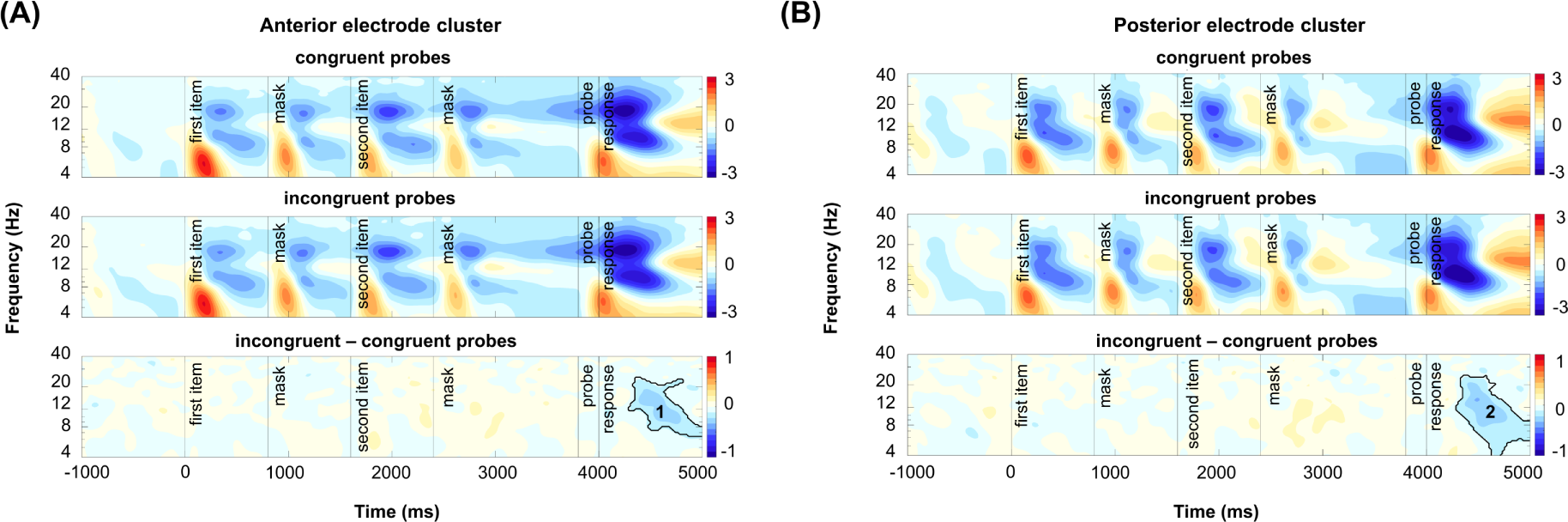
Time-frequency modulations and cluster-based permutation test results for incongruent vs. congruent probe contrast. *Note*. (A) Power was averaged across the midcentral electrode cluster for congruent and incongruent trials (upper and middle rows). Differences between the two conditions were tested with cluster-based permutation tests (lower row); (B) Power was averaged across the parieto-occipital electrodes for congruent and incongruent trials (upper and middle rows). Differences between the two conditions were tested with cluster-based permutation tests (lower row). Solid lines illustrate significant clusters with a t-mass value smaller than 1st or larger than the 99th percentile of the distribution of significance probabilities of the null distribution.

## Discussion

Previous research investigating the concurrent storage of auditory and visual features in working memory has predominantly adopted a dual-task approach, considering the two modalities as competitors for a shared underlying resource. Here, we introduce a novel multisensory delay-match-to-sample paradigm, in which bottom-up integration of audio-visual features is promoted by spatiotemporal synchrony. The paradigm included three variants of task-instructions: a cross-modal conjunction condition, in which participants needed to maintain audio-visual objects, as well as two single-feature conditions, in which participants attended to visual or auditory features, respectively. The present study reveals three major findings: First, we show that when attending to only one modality dimension of a compound audio-visual object, task-irrelevant features are encoded into working memory, as indexed by interference with behavioral performance. Although ERP results revealed that the task-irrelevant features of attended objects are not actively maintained in working memory, they tax attentional resources, as shown by alpha power dynamics during recall. Secondly, our behavioral results corroborate the notion that audio-visual working memory storage is both feature- and object-based. Finally, we provide novel insights into the oscillatory dynamics underlying audio-visual working memory. We show that attentional resources, supporting linking between auditory and visual modalities, are specifically recruited at recall rather than during maintenance or encoding. In the following, we discuss how the current findings advance our understanding of multisensory interactions in working memory.

### Are task-irrelevant cross-modal features automatically encoded and stored in working memory?

In the present study, the manipulation of probe congruency allowed us to investigate whether task-irrelevant features of an audio-visual object are encoded into working memory, when participants attend to only one of the two concurrently present modalities. Previous work had shown that when a unimodal object is attended, attention spreads towards a task-irrelevant but simultaneously occurring object from a different modality (Busse et al., 2005; Donohue et al., 2011). Critically, here we show for the first time that the effect of this cross-modal spread of attention during encoding persists throughout the maintenance interval of an audiovisual working memory task. This was indexed by interference of task-irrelevant cross-modal features at recall, suggesting that the task-irrelevant features were still represented to some degree This finding aligns with previous work in the visual working memory domain, showing that task-irrelevant features are not fully dropped from working memory (Schneider et al., 2015, 2016), but can still be retrieved, albeit only with low precision (Shin & Ma, 2016), as well as with evidence, demonstrating that task-irrelevant visual features can be decoded based on the multivariate EEG signal (Chen et al., 2021; but see also Yu & Shim, 2017, Bocincova et al., 2017). Notably, a very similar behavioral interference effect was reported in a purely auditory working memory study by Joseph and colleagues (2015), in which participants maintained sequences of whole auditory objects or were instructed to memorize only one of their component features (i.e., a spectral feature or a temporal feature). The authors found that performance was reduced when the task-irrelevant dimension of the memory objects and the probe varied randomly compared to when it was held constant. They attribute this effect to the interference of the irrelevant with the relevant dimension as well as a feature extract cost. Importantly, the authors emphasize that interference could occur both at encoding, where interference of the irrelevant dimension may add noise to the memory representation of the task-relevant feature, as well as at recall, where participants need to focus on the probe’s relevant dimension while ignoring the task-irrelevant dimension. While we cannot fully exclude that the congruency effect in the present study is partially due to interference already at encoding, the observed alpha power dynamics at recall for congruent as compared to incongruent probes, provide evidence for interference at recall. Specifically, a stronger suppression of alpha power was evident during recall when participants respond to an incongruent as compared to a congruent probe, reflecting greater attentional demands when handling incongruency. Here, for a congruent probe (i.e., both the task-relevant and the task-irrelevant probe features suggest the same response), a working memory representation containing both visual and auditory features, can serve as a template to successfully reject or accept the probe object as a whole without causing any interference. In contrast, when confronted with an incongruent probe, where processing the task-irrelevant modality would result in the selection of a false response (e.g., task-relevant feature dimension suggests a ‘yes’ response whereas the task-irrelevant feature dimension suggests a ‘no’ response), it requires additional attentional resources to selectively compare only the task-relevant features of the probe to the contents of working memory.

Notably, despite behavioral evidence for the persistence of a memory trace reflecting the task-irrelevant cross-modal feature, an ERP analysis did not provide any evidence for the active maintenance of auditory features in the attend-visual condition. Critically, Bayesian statistics provided evidence for the lack of a load-modulation of auditory SAN amplitudes in the visual condition; In contrast, in the auditory and the conjunction condition, where auditory features were task-relevant, SAN amplitudes increased with memory load, in line with previous studies, establishing the SAN as an indicator of auditory working memory storage (Alunni-Menichini et al., 2014; Guimond et al., 2011). The apparent contradiction between the behavioral and ERP results can be reconciled by models which posit that working memory content can be maintained by neural activity in an ‘activity-silent mode’ (Stokes, 2015). However, it should also be noted that we were not able to find the hypothesized memory load modulations of posterior NSW amplitudes, as an index of visual working memory storage, irrespective of whether visual information was task-relevant or irrelevant. This could be due to some prominent differences in our study design compared to previous studies, exploiting NSW ERP effects. The majority of previous studies (Diaz et al., 2021; Feldmann-Wüstefeld, 2021; Fukuda et al., 2015) used concurrent stimulus presentation, as compared to sequential presentation in the present study, and higher set sizes (up to set size 8). Finally, it remains possible that integrated audio-visual working memory items undergo some transformation which is not fully captured by ERP correlates for unimodal working memory storage.

### Are audiovisual features stored as an integrated multisensory object?

Overall, the evidence discussed so far is in line with the assumption that audio-visual features are automatically integrated in a bottom-up fashion, under conditions of audio-visual synchrony and low perceptual load (reviewed by Talsma et al., 2010) and subsequently maintained in a potentially activity-silent format (Stokes, 2015) or outside the focus of attention (Oberauer, 2002; Oberauer & Lin, 2017) . Although, on a cautionary note, it should be mentioned that the present study did now allow for an assessment of gamma power (reviewed by Engel et al., 2012) or ERP modulations according to an additive model (Besle et al., 2004) as explicit evidence for multisensory integration during encoding. Thus, despite clear evidence that both auditory and visual features are stored in working memory, in at least a residual form, in the two single-feature conditions, a critical question that remains to be addressed is whether they are stored in an integrated format or rather in separate stores. We postulate that in the present paradigm, an audio-visual representation is formed at encoding, irrespective of the task instructions. That is because we explicitly designed the task to facilitate bottom-up encoding. In accordance with the notion of a hierarchical working memory structure (Brady et al., 2011), we assume that this representation contains both a feature- and an object level, allowing participants to focus limited working memory resources on the maintenance of the feature-level in the single-feature conditions, while relying more strongly on the (integrated) object level in the conjunction condition. Corroborating the assumption of a hierarchical working memory representation on the audio-visual level (Li et al., 2022), the behavioral analysis of set size effects showed evidence for both object-based as well as feature-based storage. Critically, in line with the above model, we show that the congruency effect (i.e., reduced accuracy in incongruent compared to congruent trials) in the conjunction condition is considerably larger compared to the single-feature conditions. This suggests that focusing on audio-visual objects in the conjunction condition might come with a cost of “breaking up” this integrated template in case of an incongruent probe type. On the contrary, disentangling the congruency effect within conditions shows that correct responses to incongruent probes in the conjunction condition were faster compared to responses to congruent probes. Thus, in correct trials, switching to the feature-dimension of the representation appeared to not come at a cost. On the neural level, we do not find conclusive evidence for the maintenance of an integrated working memory representation. The oscillatory dynamics (see also section 4.3) during the maintenance period rather suggest domain-specific modulations, while integration across modalities appears to occur specifically during recall. This could be more broadly in line with the distinction between cross-modal *binding* as a purely stimulus-driven, perceptual process and multisensory *integration* at the level of decision making, made by Bizley and colleagues (2016). Ultimately, a conclusive answer to the question whether audio-visual working memory storage relies on an integrated representation warrants further investigation.

### Oscillatory dynamics underlying audio-visual working memory

An additional central finding concerns the underlying oscillatory signatures of multisensory working memory. While previous studies have primarily focused on the neurocognitive mechanisms supporting the prioritization of one modality over the other (e.g., Spitzer & Blankenburg, 2012; van Ede et al., 2017), the present study allowed us to track oscillatory mechanisms involved in audiovisual working memory. Here, by contrasting the single-feature conditions with the conjunction condition, we adopted the following logic: Any mechanism reflecting the linking of information across modalities should be consistently evident for both comparisons of interests (auditory vs. conjunction; visual vs. conjunction). This was only the case in the recall phase, where we coherently observed stronger alpha-beta power suppression over frontal and parieto-occipital cortex in the conjunction condition compared to both single-feature conditions. That is, when the task requires an audio-visual judgement, additional attentional resources seem to be recruited via the fronto-parietal attentional network specifically at recall, rather than during maintenance or encoding. This extends previous reports of stronger alpha- and beta-power suppression accompanying recognition judgments in a cross-modal delayed-match-to-sample paradigm, requiring participants to compare a sample angle encoded in one modality (i.e., a sample angle either encoded in visual or kinesthetic modality) to a probe angle in either the same or a different modality (Seemüller et al., 2012).

Further, in the late encoding period and during maintenance, we observed differential modulations of oscillatory power, most prominent in the alpha-band, but extending into the adjacent theta and lower beta-band. Essentially, as the prioritization shifts from a focus on purely visual features to purely auditory features, with the conjunction condition residing in-between, we observe a stronger relative increase in alpha power over posterior scalp sites (see Fig 7B). In line with the postulated inhibitory function of alpha oscillations (Jensen & Mazaheri, 2010; Klimesch et al., 2011) this pattern of results aligns well with the notion of a progressive disengagement of visual areas as participants have to re-distribute attentional resources between modalities (i.e., conjunction condition) or focus exclusively on auditory features (i.e., auditory condition). Notably, we also observed a very similar pattern of results at a fronto-central cluster of electrodes (see Fig 7D). Although these electrodes were selected based on a task-independent localizer procedure, aiming at identifying electrodes that were maximally sensitive to auditory stimuli, due to methodological constraints and anatomical characteristics of auditory cortex (Clements et al., 2023), it is unlikely that the observed power modulations reflect auditory oscillations. However, consistent with our results, modulations of fronto-central alpha power have been previously observed in a cross-modal matching paradigm. Specifically, Misselhorn and colleagues (2019) observed a relative increase in power for audio-visual compared to visual-tactile attention. The authors postulate that frontal alpha oscillations reflect the origin of top-down control.

More generally, to appreciate the novelty of the present findings, it is important to again highlight that despite the ample number of studies investigating neural oscillations in the context of multisensory processing and attention on a perceptual level (for a review see (Keil & Senkowski, 2018), to the authors’ best knowledge this is the first study that explicitly exploits oscillatory mechanisms in an audio-visual working memory paradigm, with a specific focus on interactions of multisensory integration and multisensory (rather than inter-sensory) attention in working memory.

## Conclusion

Exploiting both behavioral as well as EEG data, the present study provides novel insights on interactions of attention and multisensory processing in working memory. Consistent with previous work (Busse et al., 2005; Donohue et al., 2011), we show that during encoding, attention spreads to task-irrelevant features of an audio-visual compound object, resulting in their automatic encoding into working memory. Critically, this study shows for the first time that the effects of this cross-modal spread of attention persist throughout the maintenance period of an audio-visual working memory paradigm, albeit electrophysiological results suggest that task-irrelevant features are not actively maintained. Finally, the observed oscillatory dynamics, particularly in the alpha-band, show that attentional resources underlying the linking between sensory modalities specifically occurs at the retrieval. Further research will be required to conclusively say whether audio-visual working memory storage relies on integrated multisensory object representations.

## Footnotes

1 The behavioral data to obtain this a-priori effect size measure was kindly provided to us by the authors.

